# Disrupted basal ganglia—thalamocortical loops in focal to bilateral tonic-clonic seizures

**DOI:** 10.1101/618215

**Authors:** Xiaosong He, Ganne Chaitanya, Burcu Asma, Lorenzo Caciagli, Danielle S. Bassett, Joseph I. Tracy, Michael R. Sperling

## Abstract

Focal to bilateral tonic-clonic seizures are associated with lower quality of life, higher risk of seizure-related injuries, increased chance of sudden unexpected death, as well as unfavorable treatment outcomes. Achieving greater understanding of its underlying circuitry offers better opportunity to control these particularly serious seizures. Towards this goal, we provide a network science perspective of the interactive pathways among basal ganglia, thalamus and the cortex, to explore the imprinting of secondary seizure generalization on the mesoscale brain network in temporal lobe epilepsy. Specifically, we parameterized the functional organization of both the thalamocortical network and the basal ganglia—thalamus network with resting-state functional magnetic resonance imaging in three groups of patients with different focal to bilateral tonic-clonic seizure histories. Using the participation coefficient to describe the pattern of thalamocortical connections among different cortical networks, we showed that, compared to patients with no previous history, those with positive histories of focal to bilateral tonic-clonic seizures, including both remote (none for over one year) and current (within the past year) histories, presented more uniform distribution patterns of thalamocortical connections in the ipsilateral medial-dorsal thalamic nuclei. As a sign of greater thalamus mediated cortico-cortical communication, this result comports with greater susceptibility to secondary seizure generalization from the epileptogenic temporal lobe to broader brain networks in these patients. Using interregional integration to characterize the functional interaction between basal ganglia and thalamus, we demonstrated that patients with current history presented increased interaction between putamen and globus pallidus internus, and decreased interaction between the latter and the thalamus, compared to the other two patient groups. Importantly, through a series of “disconnection” simulations, we showed that these changes in interactive profiles of the basal ganglia—thalamus network in the current history group mainly depended upon the direct but not the indirect basal ganglia pathway. It is intuitively plausible that such disruption in the striatum modulated tonic inhibition of the thalamus from the globus pallidus internus could lead to an under-suppressed thalamus, which in turn may account for their greater vulnerability to secondary seizure generalization. Collectively, these findings suggest that the broken balance between the basal ganglia inhibition and thalamus synchronization can inform the presence and effective control of focal to bilateral tonic-clonic seizures. The mechanistic underpinnings we uncover may shed light on the development of new treatment strategies for patients with temporal lobe epilepsy.

## Introduction

Over 70% of patients with focal epilepsies can occasionally experience focal to bilateral tonic-clonic seizures (FBTCS) (Forsgren et al., 1996). The prevalence of FBTCS is associated with lower quality of life (Yoo *et al.*, 2014), higher risk of seizure-related serious injuries (Lawn *et al.*, 2004) and SUDEP (sudden unexpected death in epilepsy) (Walczak *et al.*, 2001; Harden *et al.*, 2017), as well as unfavorable treatment outcomes (Bone *et al.*, 2012; Keller *et al.*, 2015). Accordingly, control of FBTCS is an important objective, particularly for patients with drug-resistant focal epilepsies, such as temporal lobe epilepsy (TLE). Why some patients suffer from uncontrolled FBTCS and others do not remains a mystery, and it is desirable to learn more about them and develop new treatment strategies.

Despite its earlier terminology as “secondarily generalized tonic-clonic seizures”, FBTCS are not truly generalized, but instead selective, affecting specific brain regions most intensely (Blumenfeld *et al.*, 2003; Holmes *et al.*, 2004; Schindler *et al.*, 2007). In particular, accumulating evidence has underscored the critical role of the thalamus and its interacting circuits during FBTCS. As an integrative hub in the brain (Hwang *et al.*, 2017), the thalamus projects distributed reciprocal connections to the entire cerebral cortex (Jones, 2007) and mediates communication between different brain networks (Sherman and Guillery, 2013). In the context of FBTCS, the thalamus may act as an extension of the epileptogenic network (Guye *et al.*, 2006), supporting the propagation of ictal activity to widespread cortical networks (Castro-Alamancos, 1999) by synchronizing abnormal cortical–subcortical electrical discharges (Blumenfeld, 2002; Norden and Blumenfeld, 2002). For example, thalamic hyperactivity has been observed during FBTCS (Hamandi *et al.*, 2006; Blumenfeld *et al.*, 2009). Compared to patients with focal seizures only, patients with additional FBTCS present extra thalamic atrophy (Yang *et al.*, 2017) and disrupted thalamocortical connections both structurally (Keller *et al.*, 2015; Chen *et al.*, 2019) and functionally (He *et al.*, 2015; Peng and Hsin, 2017).

As a “braking system” between the cortex and thalamus (Vuong and Devergnas, 2018), the basal ganglia (BG) appear to be involved in FBTCS as well. The BG are a complex group of nuclei, including the striatum (putamen, caudate, and ventral striatum), globus pallidus (externus, GPe, and internus, GPi), subthalamic nucleus (STN), and substantia nigra (SN) (Wichmann and DeLong, 2012). The BG act in a topographically segregated manner interacting with thalamus and cortex, constituting several parallel circuits including the direct (cortex—striatum—GPi—thalamus—cortex) and indirect (cortex—striatum—GPe—STN—GPi—thalamus—cortex) pathways (Alexander *et al.*, 1986, 1990; Smith *et al.*, 1998). While the BG are also hyperactive during FBTCS (Blumenfeld *et al.*, 2009), and patients with FBTCS also present additional BG atrophy compared to those without (Yang *et al.*, 2017), the BG may play an anticonvulsive role during seizures (Rektor *et al.*, 2012). A recent intracranial EEG study has reported changes in cortex—striatum synchronization level throughout focal seizures as a part of an endogenous mechanism controlling the duration and termination of abnormal oscillations (Aupy *et al.*, 2019). In some reports, the occurrence of dystonia, a semiology associated with increased BG activity (Cooper, 1962; Mizobuchi *et al.*, 2004), is negatively correlated with the presence of FBTCS in TLE (Cleto Dal-Cól *et al.*, 2008; Feddersen *et al.*, 2012; Popovic *et al.*, 2012; Uchida *et al.*, 2013). Specifically, Rektor *et al.* (2002, 2011) found the BG are only involved when the ictal activity has spread to other cortical areas, *e.g.*, during secondary seizure generalization.

To date, the mechanisms implicated in such an inhibitory role for the BG remain largely hypothetical. In TLE, the BG-thalamocortical loops have been implicated in studies involving animal models [see a recent review in (Vuong and Devergnas, 2018)]. Yet, in *vivo*, the organization of these circuits, *n.b.*, the interaction between the BG and the thalamus, have rarely been studied, particularly with regard to FBTCS. Given that the thalamus has long been recognized as a seizure synchronizer (Guye *et al.*, 2006; Bertram *et al.*, 2008), we suspect that the presence of FBTCS may reflect a broken balance between BG inhibition and thalamic synchronization, which could reshape the interactions along the BG-thalamocortical loops.

To test this intuition, we employed resting-state functional MRI (rsfMRI) to assess the associations between the BG–thalamus–cortex interactions and the prevalence of FBTCS. Benefiting from the emerging computational tools and conceptual frameworks of network neuroscience (Bassett and Sporns, 2017), we evaluated the topological organization of these networks. First, using the participation coefficient, we assessed the degree to which the thalamus played the role of a connector hub in thalamocortical networks, enhancing its ability to facilitate communication among brain networks (Hwang *et al.*, 2017). We expected that greater hubness would promote cross-network communication for broader synchronization, such as that observed in secondary generalization. Second, using the interregional integration (Bassett *et al.*, 2015), we assessed the interactions within the BG–thalamus circuitry. We predicted that the occurrence of FBTCS would trigger the proposed BG inhibitory mechanism more excessively and eventually reshape the network’s organization.

Importantly, both the BG and the thalamus are involved not only in FBTCS, but also in more restricted focal seizures (Blumenfeld *et al.*, 2004), and they contribute to the disruption of large-scale cortico-subcortical functional networks in focal epilepsies (Výtvarová *et al.*, 2017). To address the unique network characteristics associated with FBTCS, our main comparisons were made across three groups of TLE patients with distinct histories with respect to FBTCS (*i.e.*, *none*, *remote*, and *current*; for definitions, see *Methods*). We hypothesized that the topological characteristics of thalamocortical and BG–thalamus networks may inform not only the presence but also the effective control of FBTCS.

## Methods

### Participants

Ninety-six patients with refractory unilateral TLE were recruited from the Thomas Jefferson Comprehensive Epilepsy Center. Diagnosis was determined by a multimodal evaluation including neurological history and examination, interictal and ictal scalp video-EEG, MRI, PET, and neuropsychological testing (Sperling, 1996). As previously published, localization was determined after determining that the testing was concordant for unilateral temporal lobe epilepsy. Patients were excluded from the study for any one of the following reasons: previous brain surgery, evidence for extra-temporal or multifocal epilepsy by history or testing, contraindications to MRI, or hospitalization for any Axis I disorder listed in the DSM-5 (Diagnostic and Statistical Manual of Mental Disorders, V). Depressive disorders were admissible given the high comorbidity of depression and epilepsy (Tracy *et al.*, 2007).

All patients had focal impaired awareness seizures (FIAS) and/or FAS (focal aware seizures), and some had FBTCS as well. For the purpose of this study, patients were placed into one of three groups (32 participants each) based on their history at the time of scanning: (1) patients who had never had any FBTCS events during their lifetime were assigned to the *none-FBTCS* group; (2) patients who had a remote history of FBTCS, but none for one year or more prior to scanning, were assigned to the *remote-FBTCS* group; and (3) patients who had recurrent FBTCS within one year prior to scanning were assigned to the *current-FBTCS* group. To provide a neuroimaging reference, 32 demographically-matched healthy controls were also recruited (*Supplementary Table S3*). All controls were free of psychiatric or neurological disorders based on health screening measures. This study was approved by the Institutional Review Board for Research with Human Subjects at Thomas Jefferson University. All participants provided informed consent in writing.

### Imaging Acquisition and Preprocessing

Onsite MRI data, including T1-weighted structural image and 5-minute rsfMRI scan, were obtained from all participants. Details regarding MRI acquisition and data preprocessing, including mitigation of motion artifact with spike regression and scrubbing (Satterthwaite *et al.*, 2013), are described in *Supplementary Methods*.

### Functional Parcellation of Striatum and Thalamus

We employed a hybrid method to define regions of interest (ROIs) using both structural and functional information. Components of the BG were structurally defined with the “atlas of the basal ganglia” (ATAG, https://www.nitrc.org/projects/atag/) (Keuken *et al.*, 2014), including the striatum, globus pallidus externus (GPe) and internus (GPi), subthalamic nucleus (STN), and substantia nigra (SN). We also localized the spatial extent of the thalamus based on the Morel atlas (Morel *et al.*, 1997; Krauth *et al.*, 2010). Due to the relatively small size of the GPi, GPe, STN, and SN (*Supplementary Table S1*), these components were treated as ROIs without any additional processing. For the striatum and thalamus, we then performed a masked independent component analysis (ICA) based functional parcellation (Moher Alsady *et al.*, 2016) to divide them into functionally distinctive subdivisions. Although these structures could be further broken down with anatomical information, ICA provided specific functional dissociations between the subdivisions, which can be crucial for studying their functional interactions. To ensure that this functional parcellation remains neutral across all experimental groups, the procedure was performed on an independent set of preprocessed rsfMRI data from the Human Connectome Project (HCP) (https://www.humanconnectome.org/). Descriptions regarding this HCP dataset are provided in *Supplementary Methods*.

Briefly, we masked the preprocessed HCP data with unilateral striatum and thalamus ROIs and spatially smoothed the data with a 6 mm kernel. We then performed ICA to generate a functional parcellation by applying a “winner-take-all” strategy (Moher Alsady *et al.*, 2016). To determine how many subdivisions were optimal for the unilateral striatum and thalamus, we ran the masked ICA across several dimensionalities for each mask using a split-half cross-validation strategy (Moher Alsady *et al.*, 2016). We found that a dimensionality of 10 for bilateral striatum and a dimensionality of 7 for bilateral thalamus provided the best cross-validation stability (*Supplementary Figure 1*). Accordingly, we parceled each striatum into 10 subdivisions and each thalamus into 7 subdivisions, which were generally symmetric in location and size (Figure 1A).

**Figure 1.**
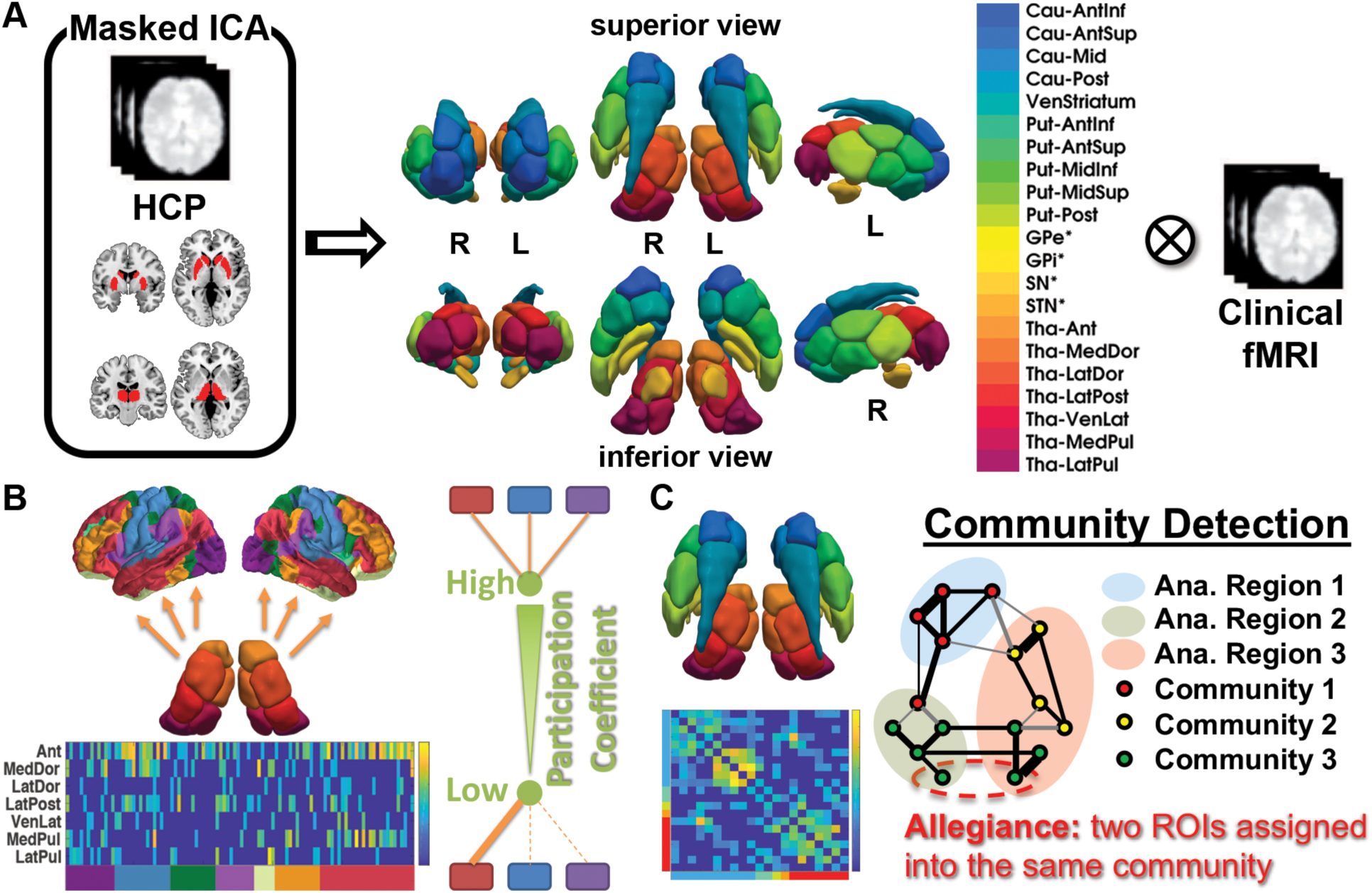
Schematic overview of the analytical pipeline. (*A*) We applied masked ICA (Moher Alsady *et al*., 2016) on an independent resting-state functional MRI dataset from the Human Connectome Project (HCP, n = 100) to generate functional parcellations of the striatum (10 parcels) and thalamus (7 parcels). In addition, anatomical masks for globus pallidus externus (GPe) and internus (GPi), subthalamic nucleus (STN), and substantia nigra (SN) were directly adopted from the ATAG atlas (Keuken *et al*., 2014), yielding a final parcellation scheme of the basal ganglia—thalamus network with 21 regions of interest (ROIs) per hemisphere. These ROIs were then used to extract time-series from the clinical resting-state functional MRI data collected in this study. (*B*) Based on the Schaefer Atlas (Schaefer *et al*., 2018), we estimated thalamocortical functional connectivity between each thalamic parcel and cortical ROIs, and then sorted them by 7 predefined resting-state networks (Yeo *et al*., 2011). We used the participation coefficient to represent the distribution pattern of thalamocortical functional connections across different resting-state networks. The more uniform the distribution, the higher the participation coefficient, and *vice versa*. (*C*) We also estimated the functional connectivity matrix of the basal ganglia—thalamus network, on which we applied a community detection algorithm, to identify groups of ROIs with higher preference for interacting with each other (*i.e.*, communities) (Newman and Girvan, 2004; Reichardt and Bornholdt, 2006; Blondel *et al.*, 2008). We used interregional integration to represent the probability of all the ROIs from two different anatomical origins being assigned to the same community (*i.e.*, allegiance) over iterative applications of this algorithm (specifically, 1000 optimizations of a modularity quality index).

Based on this hybrid method, 21 ROIs were generated for each hemisphere (*Supplementary Table S1*). We then extracted the mean time-series from the preprocessed onsite rsfMRI data using these subcortical ROIs, as well as 200 cortical ROIs defined in Schaefer *et al.* (2018). For the subsequent network analyses, these time-series were used to generate Pearson correlation matrices within each hemisphere, respectively, keeping in line with the unilateral nature of the BG– thalamocortical loops (Alexander *et al.*, 1986; Parent and Hazrati, 1995; Jones, 2007). We further removed the unreliable connections (*e.g.*, correlations with an FDR-corrected *P* value > 0.05) by setting their weights to zero, and took the absolute value of all remaining connection weights for all the matrices.

### Distribution Pattern of Thalamocortical Connections

To test our hypothesis that the thalamus plays a role as a connector hub in the brain to facilitate secondary generalization of seizures, we estimated the participation coefficient (PC) of each thalamic parcel to describe how uniformly the intrinsic functional connectivity (FC) between the thalamic and the cortical ROIs were distributed across seven well-known cortical resting-state networks (Yeo *et al.*, 2011) (see details in *Supplementary Methods, Figure 1B*). In a nutshell, a thalamic parcel with more uniformly distributed thalamocortical connections will present a PC closer to 1, and in contrast, one with more varyingly distributed connections will present a PC closer to 0.

### Community Detection based Interregional Integration

To investigate the interaction between the BG and the thalamus, we applied a generalized Louvain-like community detection algorithm (Newman and Girvan, 2004; Reichardt and Bornholdt, 2006; Blondel *et al.*, 2008) on the FC matrix among the 21 subcortical BG and thalamic ROIs, to identify groups of ROIs with higher preference for interacting with each other (*i.e.*, communities). This algorithm is described in *Supplementary Methods*. We used interregional integration to represent the probability of all the ROIs from two different partitions being assigned to the same community over iterative applications of the algorithm (see *Supplementary Methods*). Intuitively, a higher value of integration represented a higher probability of the members from one partition being assigned to the same community with the members from another partition, potentially suggesting a higher functional interaction between these two anatomical structures (Figure 1C).

For each BG–thalamus network, the 21 ROIs were grouped following their main anatomical boundaries into five partitions: the striatum (n=10), GP (n=2), STN (n=1), SN (n=1), and thalamus (n=7), yielding 10 pairwise integration values between every possible pair. Subsequently, the striatum was further broken down into caudate (n=4), ventral striatum (n=1), and putamen (n=5), while the thalamus was further broken down into anterior (n=1), medial-dorsal (n=1), lateral-ventral (n=3), and posterior (n=2) nuclear groups (Krauth *et al.*, 2010) to finely probe pairwise functional interactions. Previously, we quantitatively demonstrated that the estimated integration value is independent of partition size [*i.e.*, number of ROIs in a partition, see details in (He *et al.*, 2018)]. Any, such bias would be distributed uniformly across the groups, and hence would not affect our between-group comparisons.

To further confirm that our results were relevant to the topological organization of the BG– thalamus network, we employed a random network null model as a benchmark (Rubinov and Sporns, 2011), by testing whether interregional integration estimated from the null networks with random topological organization could produce results similar to those observed in real brain networks (see details in *Supplementary Methods*).

### Permutation-based Statistical Testing

To minimize the bias of data distribution to our statistical inferences and to correct for multiple comparisons, we implemented a permutation-based method as our main statistical strategy (Groppe *et al.*, 2011). Briefly, the observed statistic for each variable is compared to the distribution of the most extreme statistic across the entire family of tests for each possible permutation, thereby producing multiple comparison corrected *P*-values by controlling the family-wise error rate (Blair and Karniski, 1993) (details are provided in *Supplementary Methods*). We performed 1,000,000 permutations each time, and used either *t* or *F* statistics as appropriate.

Since the hemispheric lateralization of the seizure focus may be irrelevant for FBTCS history in TLE patients (Bone *et al.*, 2012), the network properties of the right TLE patients were flipped left to right, allowing all statistical analyses to be conducted in accordance with the site of ictal onset (left, ipsilateral; right, contralateral). This method has been undertaken previously to increase statistical power (Bernhardt *et al.*, 2010; He *et al.*, 2017; Yang *et al.*, 2017). Comparisons for common demographic and clinical information were made with standard parametric tests such as a one-way ANOVA or Chi-Square, conducted using IBM® SPSS® v25. The alpha level was set at *P*<0.05 for both parametric and nonparametric tests.

### Data availability

The data that support the findings of this study are available from the corresponding author upon reasonable request.

## Results

### Demographical, Clinical, and Data Quality

All three patient groups were matched by age, sex, and handedness (Oldfield, 1971), as well as the lateralization of their seizure focus (Table 1). As the presence of FIAS can also affect the BG and the thalamus (Blumenfeld *et al.*, 2004), we compared the frequency of FIAS among these patients, and found no significant differences. However, these patients did have differences in some other clinical features. Specifically, we noted the *remote*-FBTCS group showed earlier age at epilepsy onset, and longer disease duration compared to the other two groups. In addition, the *none*-FBTCS group had more varied temporal pathologies as evidenced by presurgical MRI scans. To test for group differences mainly driven by FBTCS history, the influence of these demographic and clinical variances needed to be minimized. Accordingly, we treated them as potential confounding factors and regressed them out. We further examined two main factors that may also bias the subsequent FC analyses, namely the anatomical structure where fMRI signals were extracted and the data quality regarding head motion control and community detection. Although we did not observe any significant group differences (*Supplementary Result 1, Tables S2-S4*), we sought to minimize their impact by confound regression as well.

**Table 1.**
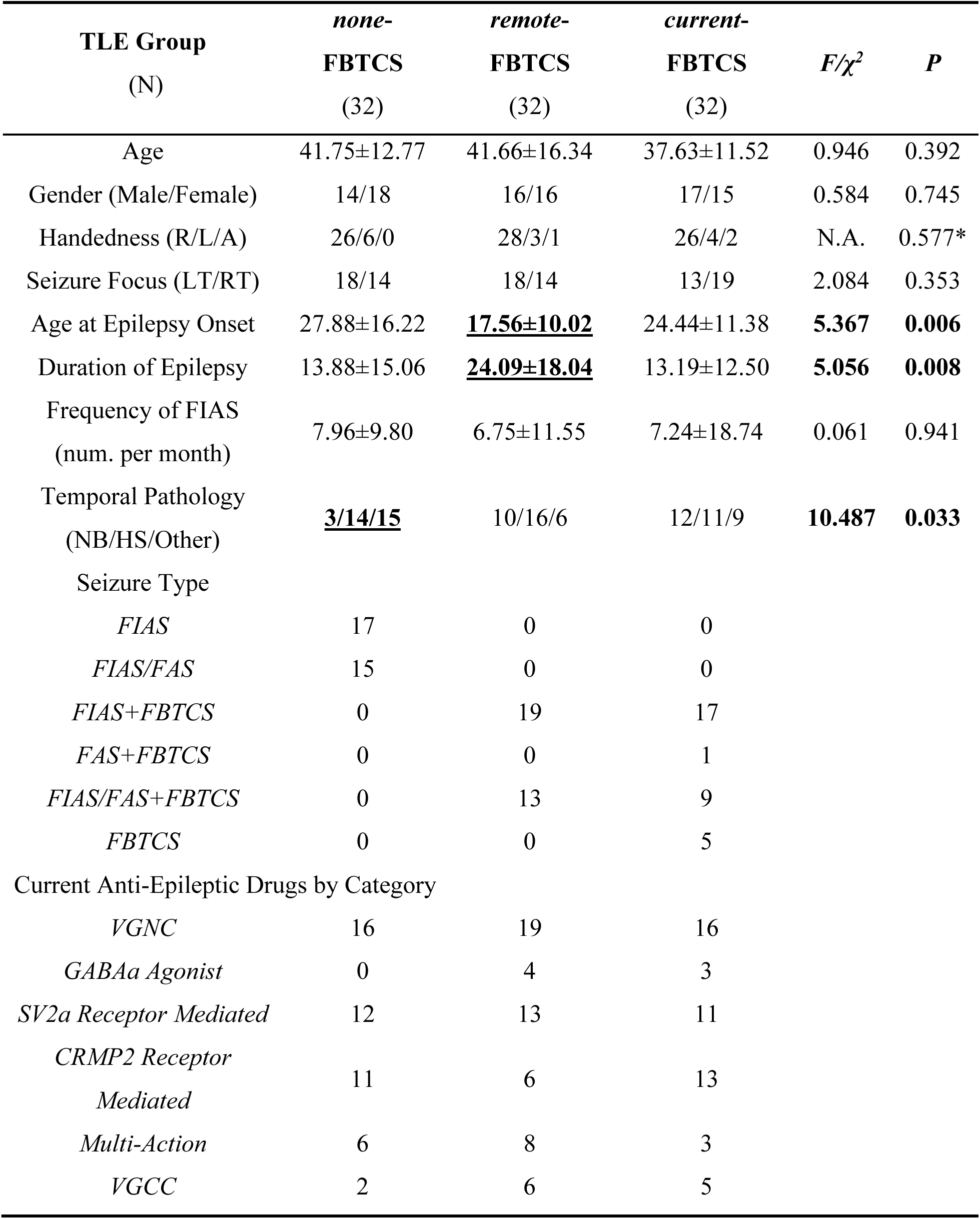
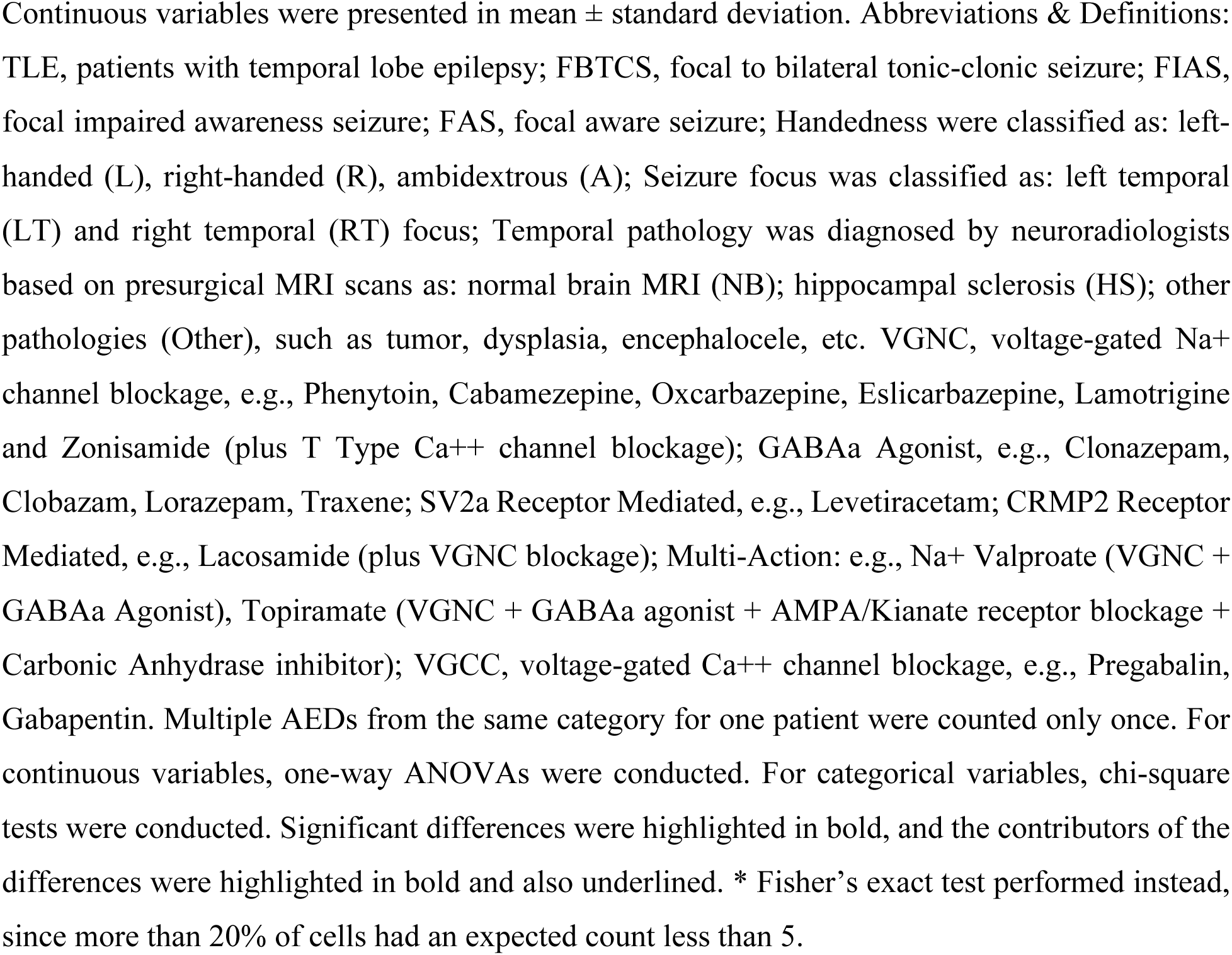
Sample demographic and clinical characteristics.

| **TLE Group**<br>**(N)** | ***none-***<br>***FBTCS***<br>**(32)** | ***remote-***<br>***FBTCS***<br>**(32)** | ***current-***<br>***FBTCS***<br>**(32)** | ***F/χ<sup>2</sup>*** | ***P*** |
| --- | --- | --- | --- | --- | --- |
| Age | 41.75±12.77 | 41.66±16.34 | 37.63±11.52 | 0.946 | 0.392 |
| Gender (Male/Female) | 14/18 | 16/16 | 17/15 | 0.584 | 0.745 |
| Handedness (R/L/A) | 26/6/0 | 28/3/1 | 26/4/2 | N.A. | 0.577* |
| Seizure Focus (LT/RT) | 18/14 | 18/14 | 13/19 | 2.084 | 0.353 |
| Age at Epilepsy Onset | 27.88±16.22 | **<u>17.56±10.02</u>** | 24.44±11.38 | **5.367** | **0.006** |
| Duration of Epilepsy | 13.88±15.06 | **<u>24.09±18.04</u>** | 13.19±12.50 | **5.056** | **0.008** |
| Frequency of FIAS<br>(num. per month) | 7.96±9.80 | 6.75±11.55 | 7.24±18.74 | 0.061 | 0.941 |
| Temporal Pathology<br>(NB/HS/Other) | **<u>3/14/15</u>** | 10/16/6 | 12/11/9 | **10.487** | **0.033** |
| Seizure Type |  |  |  |  |  |
| *FIAS* | 17 | 0 | 0 |  |  |
| *FIAS/FAS* | 15 | 0 | 0 |  |  |
| *FIAS+FBTCS* | 0 | 19 | 17 |  |  |
| *FAS+FBTCS* | 0 | 0 | 1 |  |  |
| *FIAS/FAS+FBTCS* | 0 | 13 | 9 |  |  |
| *FBTCS* | 0 | 0 | 5 |  |  |
| Current Anti-Epileptic Drugs by Category |  |  |  |  |  |
| *VGNC* | 16 | 19 | 16 |  |  |
| *GABAa Agonist* | 0 | 4 | 3 |  |  |
| *SV2a Receptor Mediated* | 12 | 13 | 11 |  |  |
| *CRMP2 Receptor*<br>*Mediated* | 11 | 6 | 13 |  |  |
| *Multi-Action* | 6 | 8 | 3 |  |  |
| *VGCC* | 2 | 6 | 5 |  |  |

In total, 12 potential confounding factors were identified, including age, sex, handedness, seizure focus side, FIAS frequency, age at epilepsy onset, epilepsy duration, MRI-evidenced temporal pathology, mean framewise displacement (Jenkinson, 2002), number of spikes scrubbed, network density, and total subcortical gray matter volumes. All confounding factors were regressed out from the network properties with a linear regression model, and the residuals were taken for subsequent statistical inferences.

### Distribution Pattern of Thalamocortical Connections

Consistent with previous work (Hwang *et al.*, 2017), we observed high PC values across all 8 thalamic nuclear groups (4 for each hemisphere) in all three patient groups. To further explore group differences in PC, we regressed out the confound variables and compared the PC across patient groups with a permutation-based *F* test. Since the thalamocortical connections were specifically tested here, the confound of total subcortical gray matter volume was replaced with thalamus volume.

After correction for multiple comparisons, a significant group difference was only found in the ipsilateral medial-dorsal nuclei [*F*_(2,93)_=8.202, *P*_corr_=0.003], where the *none*-FBTCS group showed lower PC compared to the other two groups (*none*<*remote*: *P*_Bonferroni_=0.001; *none*<*current*: *P*_Bonferroni_=0.004, Figure 2A). This result can be reproduced without confound regression, with PC estimated from binary matrices, or when bilateral functional thalamocortical connections were also taken into account (*Supplementary Result 2*).

**Figure 2.**
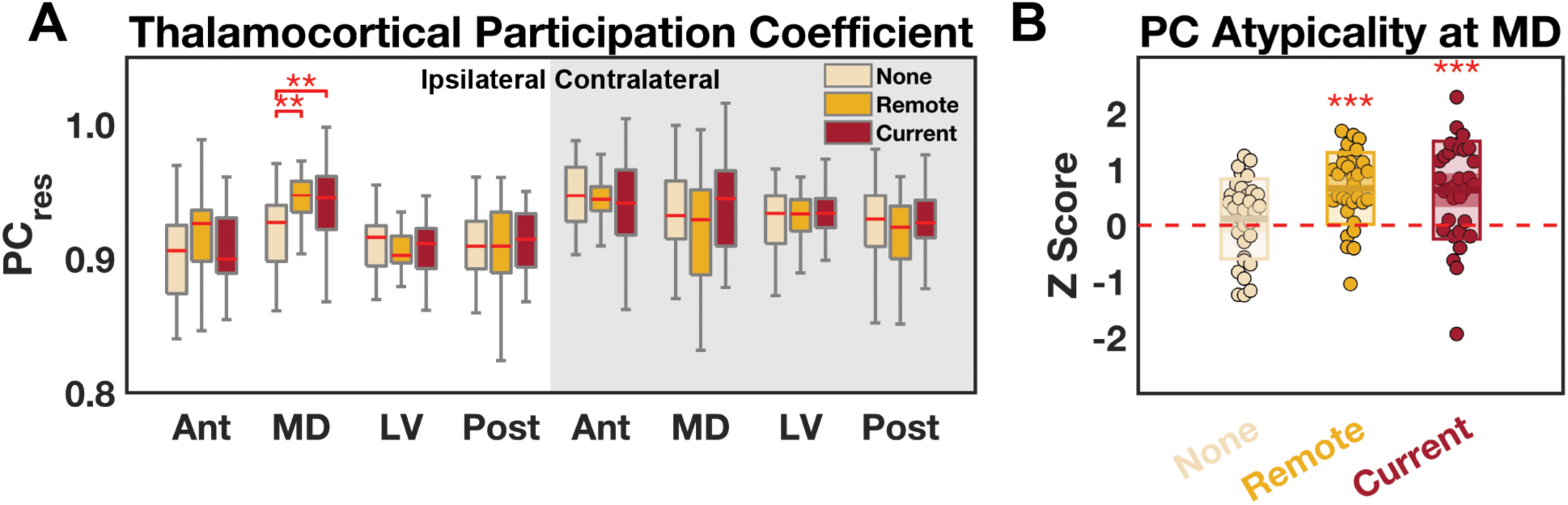
Thalamocortical participation coefficients compared across the three patient groups. (*A*) A significant difference was found in the ipsilateral medial-dorsal (MD) thalamic nuclear group. (*B*) Atypicality was estimated in reference to data obtained from a matched healthy control group. The *remote*- and *current*-FBTCS groups presented Z scores significantly higher than 0, but not the *none*-FBTCS group. The full spectrum of Z scores and data from healthy control group are presented in *Supplementary Figure 2*. Res: residual after confound regression; Ant, anterior, MD, medial-dorsal, LV, lateral-ventral, Post, posterior thalamic nuclear groups. **, *P* < 0.01; ***, *P* < 0.001. Statistics were obtained via a non-parametric permutation test controlling for multiple comparisons. The central mark indicates the median, and the bottom and top edges of the box indicate the 25^th^ and 75^th^ percentiles, respectively.

To determine the atypicality in the PC patterns, we brought in a group of demographically-matched healthy controls to provide a neuroimaging baseline. The five clinical confounding factors were excluded from this analysis because they did not apply to healthy controls. In addition, as there was no way to flip healthy controls’ data to match the ipsilateral versus contralateral side in right TLE patients, we instead calculated the deviation score of PC [*Z_pat_* = (*PC_pat_* − *μ_con_*)/*σ_con_*, where *μ_con_* and *σ_con_* were the mean and standard deviation of the same thalamic PC from the healthy controls] for each patient at each hemisphere and flipped the Z score of right TLE afterwards. We used a permutation-based one-sample *t*-test to assess the Z scores of the medial-dorsal nuclei, and found the *none*-FBTCS group presented Z scores comparable to 0 (*t*_31_=0.892, *P*_corr_=0.758), while the *remote*-FBTCS group (*t*_31_=5.814, *P*_corr_=1.6×10^-5^) and *current*-FBTCS group (*t*_31_=4.061, *P*_corr_=0.001) presented Z scores significantly higher than 0 (Figure 2B). These findings suggest that the latter two groups had elevated PC values in comparison to healthy controls.

### Interregional Integration between Basal Ganglia and Thalamus

To determine whether the interregional integration between the BG and the thalamus differed among patients with different FBTCS history, we applied the permutation-based *F* test to compare the 20 pairwise integration measures (10 for each hemisphere) across the three patient groups, after regressing out all confound variables. After correction for multiple comparisons, significant group differences were found in the striatum–GP integration [*F*_(2,93)_=7.693, *P*_corr_=0.016] and the GP– thalamus integration [*F*_(2,93)_=10.446, *P*_corr_=0.002], both on the ipsilateral side. Specifically, the *current*-FBTCS group presented higher striatum–GP integration (*current*>*none*: *P*_Bonferroni_=0.002; *current*>*remote*: *P*_Bonferroni_=0.004) and lower GP–thalamus integration (*current*<*none*: *P*_Bonferroni_=1.7×10^-4^; *current*<*remote*: *P*_Bonferroni_=0.001), compared to the other two groups (Figure 3A). Results remain significant when not regressing out confounding variables, as well as when using different resolution parameter values during community detection (*Supplementary Result 3*), suggesting their robustness to variations in our analysis strategy.

**Figure 3.**
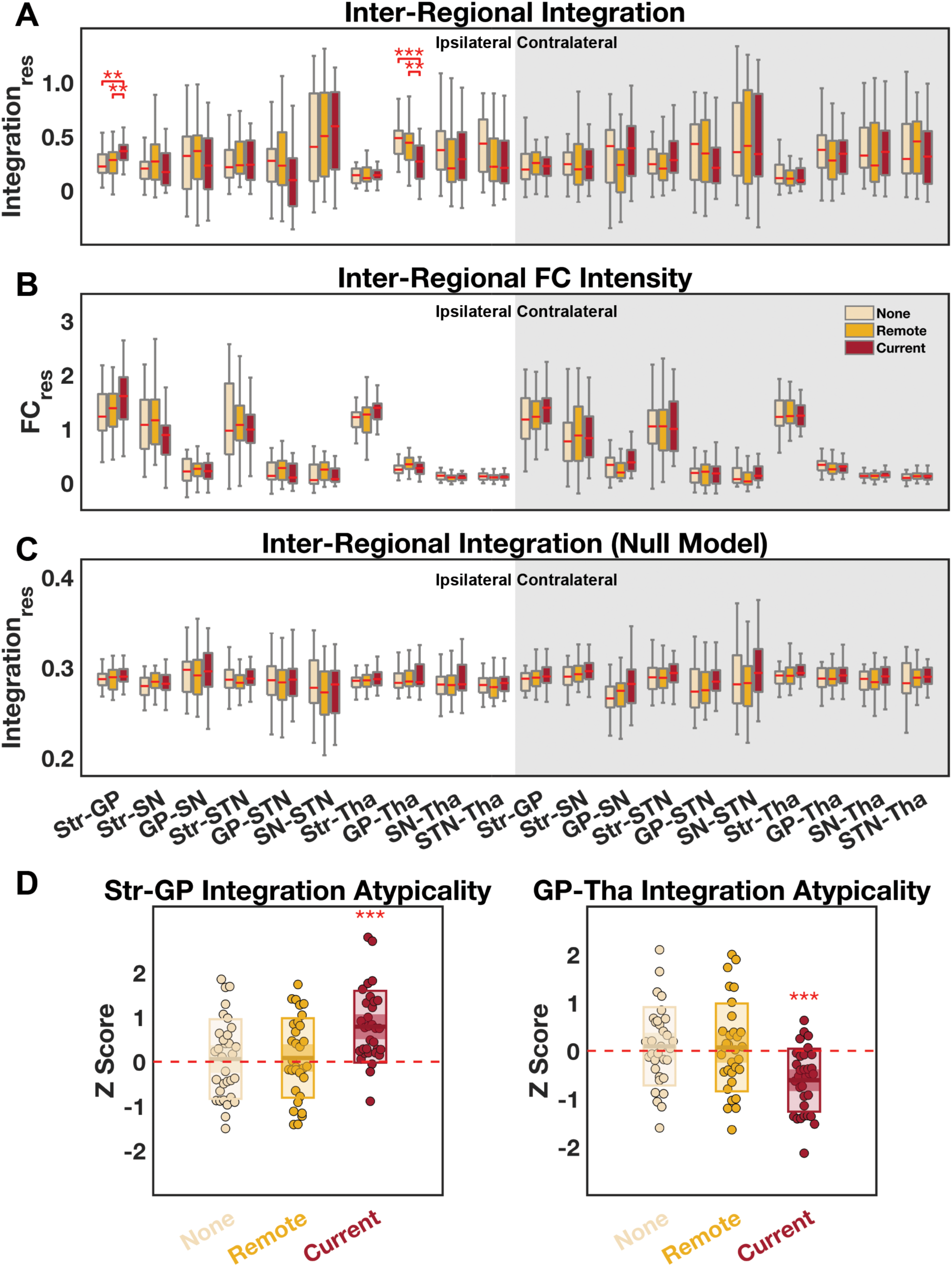
Pairwise interregional integration of the basal ganglia—thalamus network compared across the three patient groups. (*A*) Significant differences in interregional integration were found at the striatum—GP and GP—thalamus pairs. (*B*) No significant differences were found when comparing pairwise functional connectivity instead of integration estimates. (*C*) No significant differences were found in a null model in which the original functional connectivity matrix was randomly rewired, preserving both the degree and strength distributions. (*D*) Atypicality was estimated in reference to data from a matched healthy control group. The *current*-FBTCS groups presented Z scores significantly different from 0, but not the other two groups. The full spectrum of Z scores and data from healthy control group are presented in *Supplementary Figure 3*. Res: residual after confound variable regression; Str, striatum; GP, globus pallidus; SN, substantia nigra; STN, subthalamic nucleus; Tha, thalamus. **, *P* < 0.01; ***, *P* < 0.001. Statistics were inferred with a non-parametric permutation-based method controlling for multiple comparisons. The central mark indicates the median, and the bottom and top edges of the box indicate the 25^th^ and 75^th^ percentiles, respectively.

A naturalistic explanation was that such group differences in interregional integration simply reflected the differences in FC intensity between these region pairs. Therefore, similar to the manner in which we tested the integration, we tested the pairwise FC intensity over the aforementioned 20 possible combinations, but did not find any significant group differences [*F*’s_(2,93)_<4.381, *P*_corr_’s>0.252, Figure 3B]. Through linear regression, we found the FBTCS history difference can still explain a significant proportion of variance in interregional integration, even when the most relevant pairwise FC intensity estimates were included in the model (*Supplementary Result 4*). Collectively, these results suggest that the group differences in interregional integration cannot be simply explained by the pairwise FC intensity, but instead higher order topological differences are responsible as well.

To evaluate the role of topological organization in the BG–thalamus network, we performed the same analysis on the integration estimated from randomly rewired networks with degree and strength distributions preserved. Again after regressing out confound variables, we found no significant group difference [*F*’s_(2,93)_<4.395, *P*_corr_’s>0.210, Figure 3C]. This result verified the contribution of topological organization to the aforementioned group differences in interregional integration.

In line with the previous sections, we further calculated the Z scores of integration values to assess atypicality where significant differences emerged. We found the *none*- and *remote*-FBTCS groups presented Z scores comparable to 0 (|*t*_31_|’s<0.661, *P*_corr_’s>0.885), while the *current*-FBTCS group presented Z scores significantly different from 0 (striatum–GP: *t*_31_=5.519, *P*_corr_=1.9×10^-5^; GP–thalamus: *t*_31_=-5.268, *P*_corr_=1.6×10^-5^; Figure 3D). In other words, the *none*- and *remote*-FBTCS groups maintained a similar BG–thalamus network organization compared to those of healthy controls, while the *recent*-FBTCS group had specific reorganization that led to their unique striatum–GP–thalamus integration pattern.

### Globus Pallidus Centered Interregional Integration

As noted, the GP was the key structure associated with the changes in the BG–thalamus interactions in the *current*-FBTCS group. Accordingly, we further broke apart the GP itself by its anatomical boundary into external (GPe) and internal (GPi) segments, and explored their interregional integration with other components. Furthermore, we also broke down the striatum into caudate, ventral striatum and putamen, and the thalamus into anterior, medial-dorsal, lateral-ventral and posterior nuclear groups to provide a finer resolution assessment. Taking into consideration the STN and SN as well, we compared a total of 18 (9 for each hemisphere) pairwise integration measures across the three patient groups for GPe and GPi, respectively, after regressing out confound variables.

Interestingly, after correction for multiple comparisons, no significant group differences can be found among the interregional integration estimates with the GPe [*F*’s_(2,93)_<2.969, *P*_corr_’s>0.586, Figure 4A]. In contrast, significant group differences were found in the putamen–GPi integration [*F*_(2,93)_=9.926, *P*_corr_=0.003], the GPi–lateral-ventral nuclei integration [*F*_(2,93)_=7.792, *P*_corr_=0.014], and the GPi–posterior nuclei integration [*F*_(2,93)_=8.094, *P*_corr_=0.011], all on the ipsilateral side. Specifically, the *current*-FBTCS group presented higher putamen–GPi integration (*current*>*none*: *P*_Bonferroni_=3.8×10^-4^; *current*>*remote*: *P*_Bonferroni_=0.001), lower GPi–lateral-ventral nuclei integration (*current*<*none*: *P*_Bonferroni_=0.001; *current*<*remote*: *P*_Bonferroni_=0.008), and lower GPi– posterior nuclei integration (*current*<*none*: *P*_Bonferroni_=0.028; *current*<*remote*: *P*_Bonferroni_=4.6×10^-4^), compared to the other two groups (Figure 4B). Additionally, we observed negative correlations between the putamen–GPi integration and the two GPi–thalamus integration estimates (with lateral-ventral nuclei: *R*_93_=-0.415, *P*=2.9×10^-5^; with posterior nuclei: *R*_93_=-0.427, *P*=1.6×10^-5^), even after controlling for the group index. The results are still robustly observed when we do not regress out confound variables (*Supplementary Result 5*).

**Figure 4.**
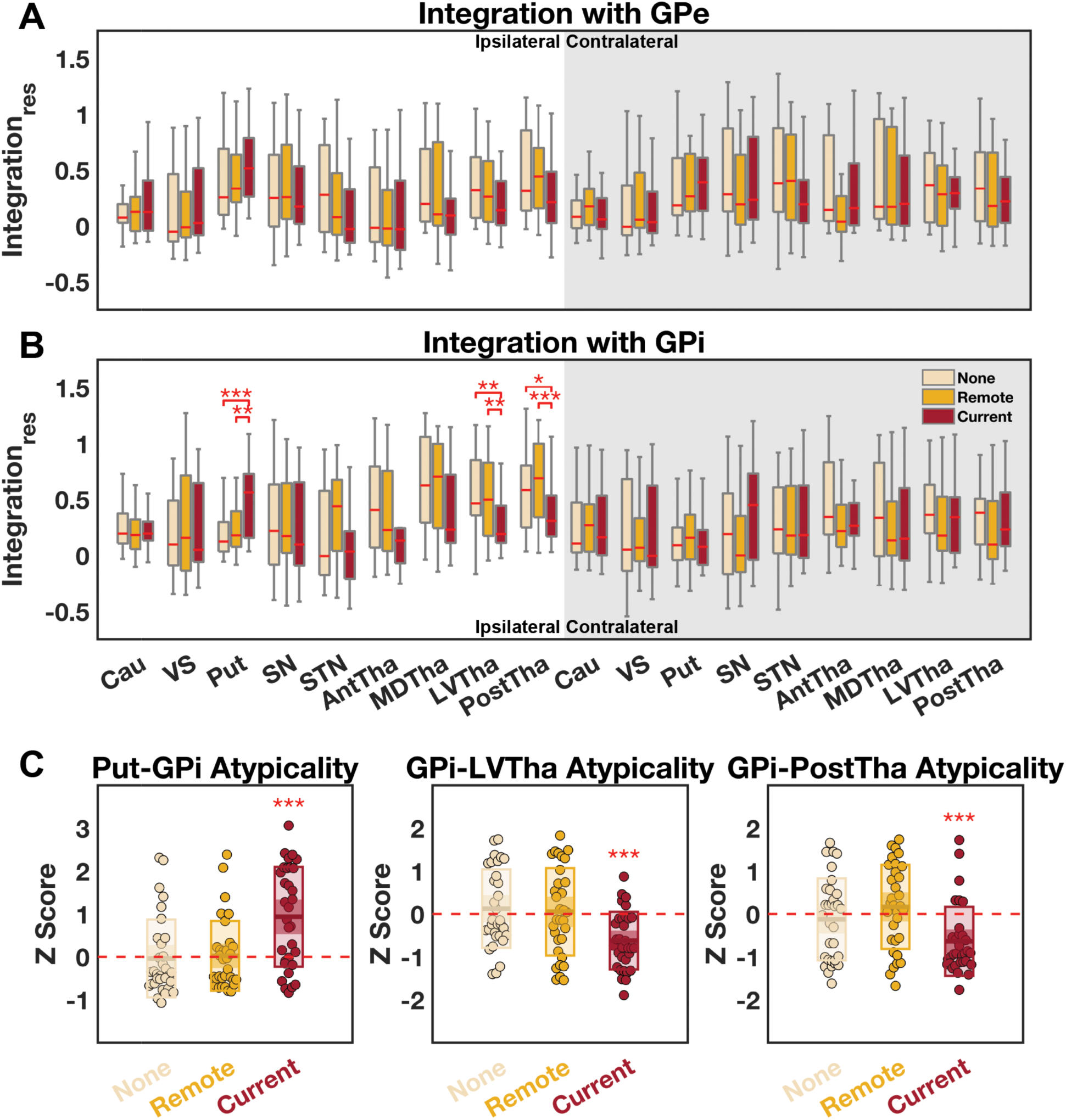
Interregional integration with GPe and GPi compared across three patient groups. (*A*) No significant differences were observed at the GPe. (*B*) Significant differences in interregional integration were observed in putamen—GPi, GPi—lateral-ventral thalamus, and GPi—posterior thalamus pairs. (*C*) Atypicality was estimated in reference to data from a matched healthy control group. The *current*-FBTCS groups presented Z scores significantly different from 0, but the other two groups did not. The full spectrum of Z scores and data from healthy control group are presented in *Supplementary Figure 4*. Res: residual after confound variable regression; GPe, globus pallidus externus; GPi, globus pallidus internus; Cau, caudate; VS, ventral striatum; Put, putamen; SN, substantia nigra; STN, subthalamic nucleus; Tha, thalamus; Ant, anterior, MD, medial-dorsal, LV, lateral-ventral, Post, posterior nuclear groups. *, *P* < 0.05, **, *P* < 0.01; ***, *P* < 0.001. Statistics were inferred with a non-parametric permutation-based method controlling for multiple comparisons. The central mark indicates the median, and the bottom and top edges of the box indicate the 25^th^ and 75^th^ percentiles, respectively.

Again, we conducted the same Z score analysis here to verify atypicality. We found the *none*- and *remote*-FBTCS groups presented Z scores comparable to 0 (|*t*_31_|’s<0.955, *P*_corr_’s>0.718), while the *current*-FBTCS group presented Z scores significantly different from 0 (putamen–GPi: *t*_31_=4.544, *P*_corr_=1.6×10^-4^; GPi–lateral-ventral nuclei: *t*_31_=-5.227, *P*_corr_=5.4×10^-5^; GPi–posterior nuclei: *t*_31_=-4.486, *P*_corr_=5.3×10^-4^; Figure 4C). These results further confirmed that the *current*-FBTCS group displayed a specific increase in putamen–GPi integration and a decrease in GPi– thalamus integration mainly with the lateral-ventral and posterior nuclei, in reference to those of healthy controls.

Theoretically, the putamen can interact with the GPi through both direct and indirect pathways. Therefore, we sought to evaluate the influence of specific FCs constituting each pathway on the observed interregional integration difference (Figure 5A), by a simulated “disconnection” method. First, we “disconnected” the direct pathway by setting the FC values of all connections between putamen and GPi to 0, and re-applied the community detection and integration estimation. After regressing out the confound variables, no significant group differences were observed among pairwise integrations with either the GPe [*F*’s_(2,93)_<2.344, *P*_corr_’s>0.798] or the GPi [*F*’s_(2,93)_<3.864, *P*_corr_’s>0.331, Figure 5B]. Specifically, if the caudate and ventral striatum (*i.e.*, the rest of the striatum) to GPi connections were zeroed out instead, the results were similar to the primary findings such that the three groups differed in pairwise integration of ipsilateral GPi [with putamen, *F*_(2,93)_=9.072, *P*_corr_=0.005, Figure 5C; with lateral-ventral nuclei, *F*_(2,93)_=6.941, *P*_corr_=0.027; with posterior nuclei, *F*_(2,93)_=7.369, *P*_corr_=0.019; others: *F*’s_(2,93)_<3.974, *P*_corr_’s>0.299] but not GPe [*F*’s_(2,93)_<3.544, *P*_corr_’s>0.407]. Second, we “disconnected” the indirect pathway by zeroing out the FC values of all connections from putamen to GPe, GPe to STN, STN to GPi, and GPe to GPi, before applying community detection and estimating integration. After regressing out confound variables, we found group differences similar to the primary findings that involved pairwise integration of ipsilateral GPi [with putamen, *F*_(2,93)_=12.730, *P*_corr_=3.2×10^-4^, Figure 5D; with lateral-ventral nuclei, *F*_(2,93)_=6.271, *P*_corr_=0.049; with posterior nuclei, *F*_(2,93)_=6.165, *P*_corr_=0.053; others: *F*’s_(2,93)_<3.871, *P*_corr_’s>0.326] but not of GPe [*F*’s_(2,93)_<5.149, *P*_corr_’s>0.111]. These results suggested that the FCs emerging from the direct, but not the indirect, pathway contribute prominently to the observed GPi integration differences.

**Figure 5.**
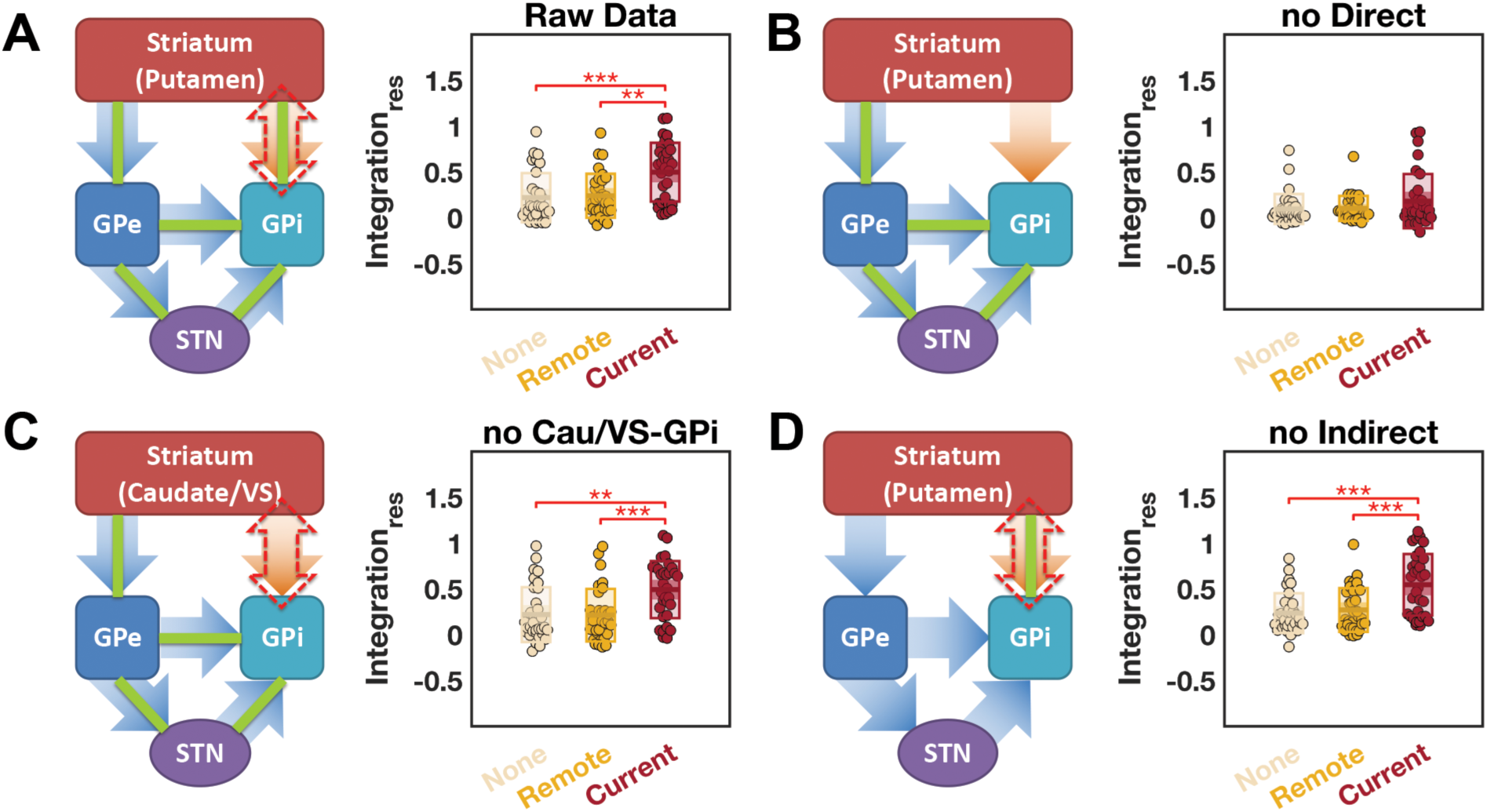
Simulated “disconnection” analyses on putamen—GPi interregional integration. To delineate specific contributions from different connections of the network, we set functional connectivity value(s) of specific connection(s) to zero (*i.e.*, “disconnected”), and then re-estimated the integration. (*A*) Before simulation, a significant group difference in putamen—GPi integration was found. (*B*) No significant putamen—GPi integration difference was found when the connections from putamen to GPi (*i.e.*, the “direct pathway”) were “disconnected”. (*C*) A significant difference in the putamen—GPi integration was found when the connections from the caudate (Cau) and ventral striatum (VS) to the GPi were “disconnected”. (*D*) A significant difference in the putamen—GPi integration was found when all connections from putamen to GPe, GPe to STN, STN to GPi, and GPe to GPi (*i.e.*, the “indirect pathway”) were “disconnected”. In the schematics, the orange arrow represents the direct pathway, the blue arrows represent both the long and short routes of the indirect pathway (Smith *et al*., 1998), the green line represents functional connectivity, and the red dashed double head arrow represents the group difference in interregional integration. GPe, globus pallidus externus; GPi, globus pallidus internus; STN, subthalamic nucleus. Bonferroni corrected Post-hoc test: **, *P* < 0.01; ***, *P* < 0.001.

We further explore the GPi–thalamus integration differences (Figure 6A) with a similar strategy. First, we “disconnected” GPi and posterior thalamic nuclei by setting all their pairwise FC values to 0 before applying community detection and estimating integration. We found that this “disconnection” simulation did not diminish the group differences in the ipsilateral GPi– thalamus integration [GPi–lateral-ventral nuclei: *F*_(2,93)_=8.390, *P*_corr_=0.009; GPi–posterior nuclei: *F*_(2,93)_=10.426, *P*_corr_=0.002, Figure 6B]. Second, we “disconnected” GPi and lateral-ventral thalamic nuclei before applying community detection and estimating integration. Although the GPi–posterior nuclei connections remained untouched, we no longer observed group differences in the ipsilateral GPi–thalamus integration [GPi–lateral-ventral nuclei: *F*_(2,93)_=4.458, *P*_corr_=0.210; GPi–posterior nuclei: *F*_(2,93)_=3.889, *P*_corr_=0.322, Figure 6C]. Third, we “disconnected” lateral-ventral and posterior thalamic nuclei instead. Interestingly this time, the group difference was only found in the ipsilateral GPi–lateral-ventral nuclei [*F*_(2,93)_=9.317, *P*_corr_=0.004] but not in the GPi– posterior nuclei [*F*_(2,93)_=1.944, *P*_corr_=0.922] integration (Figure 6D). Collectively, these results suggest that the observed GPi–posterior nuclei integration differences may be prominently contributed by FCs connecting GPi to lateral-ventral nuclei, and lateral-ventral to posterior nuclei.

**Figure 6.**
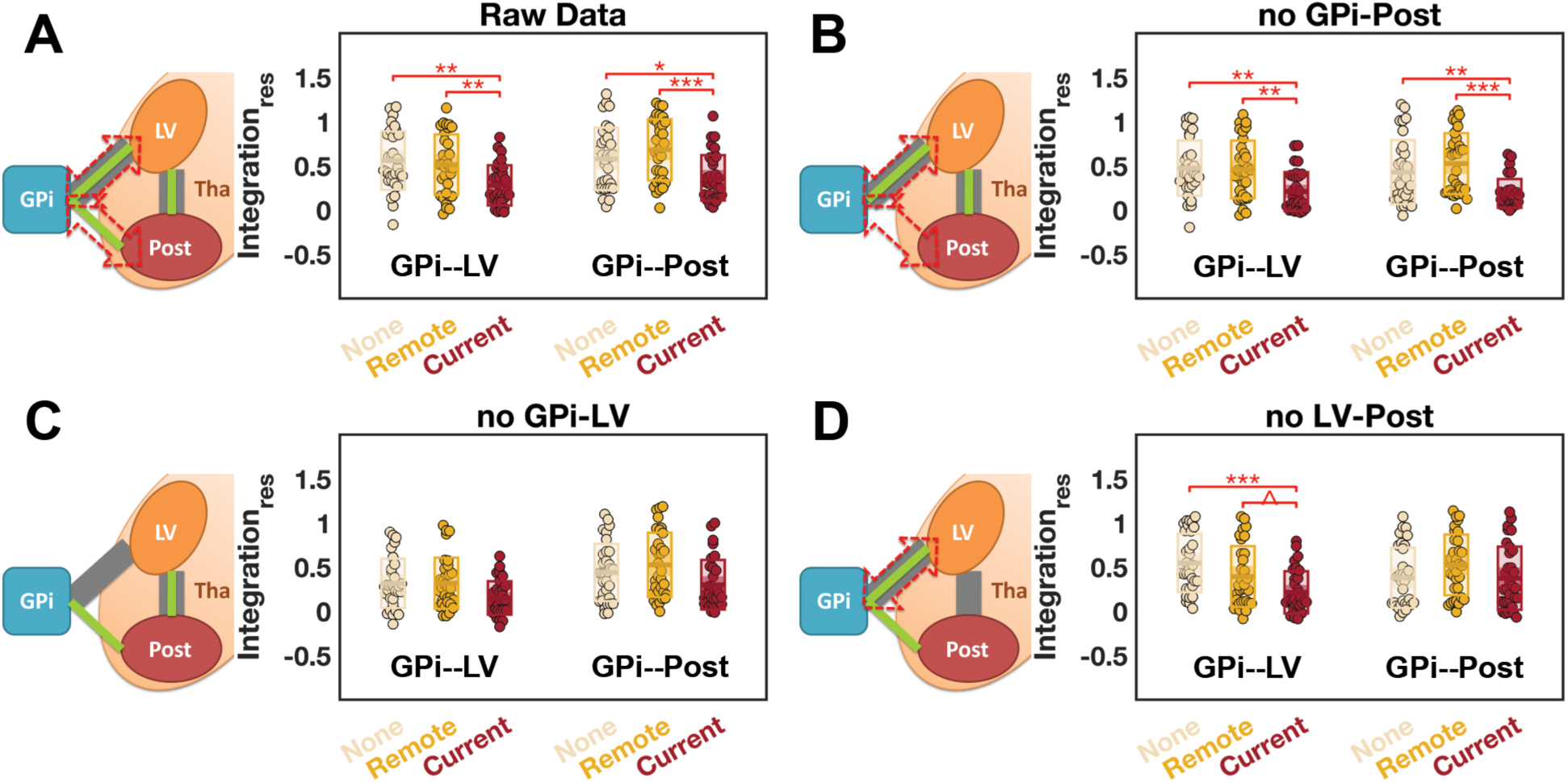
Simulated “disconnection” analyses on GPi—thalamus interregional integrations. To delineate specific contributions from different connections of the network, we set functional connectivity value(s) of specific connection(s) to zero (*i.e.*, “disconnected”), and then re-estimated the integration. (*A*) Before simulation, we observed significant group differences in GPi—lateral-ventral (LV), and in GPi—posterior (Post) thalamus integration. (*B*) Significant integration differences were still observed when the connections from GPi to the posterior thalamic nuclei were “disconnected”. (*C*) No significant integration differences were found when the connections from the GPi to the lateral-ventral thalamic nuclei were “disconnected”. (*D*) Significant integration differences were found at the GPi—lateral-ventral but not the GPi—posterior thalamus pair when all connections between the lateral-ventral and posterior thalamic nuclear groups were “disconnected”. In the schematics, thick grey lines represent anatomical connections, the green line represents functional connectivity, and the red dashed double head arrow represents the group difference in interregional integration. GPi, globus pallidus internus; Tha, thalamus; LV, lateral-ventral, Post, posterior nuclear groups. Bonferroni corrected Post-hoc test: ^, 0.05 < *P* < 0.1; *, *P* < 0.05; **, *P* < 0.01; ***, *P* < 0.001.

## Discussion

Pursuant to a network science perspective, we investigated the functional organization of the BG—thalamocortical loops in TLE patients with different FBTCS histories. We found that, compared to patients who never experienced FBTCS, patients who had a positive history of FBTCS showed more uniformly distributed thalamocortical connections, particularly in the ipsilateral medial-dorsal thalamic nuclei. Patients with uncontrolled FBTCS also presented additional atypicalities in their BG–thalamus network compared to the other two patient groups, characterized by distinct putamen–GPi–thalamus interactions. These results suggested that the topological architecture of the thalamocortical network and the BG–thalamus network together can inform both the presence and the effective control of FBTCS (Table 2).

**Table 2.** Schematic summary of findings.

| FBTCS history | Thalamus mediated<br>cortico-cortical communication | BG–thalamus interaction |
| --- | --- | --- |
| *None* | normative | normative |
| *Remote* | promoted | normative |
| *Current* | promoted | reduced |

The medial-dorsal nuclei are known to be connected to the hippocampus and other mesial temporal structures (Dolleman-Van der Weel *et al.*, 1997; Bertram and Zhang, 1999; Behrens *et al.*, 2003), and hence are heavily implicated in TLE. For instance, hypometabolism (Juhász *et al.*, 1999), structural abnormalities (Bernhardt *et al.*, 2012; Barron *et al.*, 2013), and altered thalamo-temporal connections (Keller *et al.*, 2014; He *et al.*, 2015) have been reported in the medial-dorsal nuclei of TLE patients. In particular, TLE patients with history of FBTCS presented additional atrophy (Yang *et al.*, 2017) and disruption in thalamo-temporal connections (Keller *et al.*, 2015; Chen *et al.*, 2019) in the ipsilateral medial-dorsal nuclei than those without FBTCS. The medial-dorsal nuclei respond to the onset of limbic seizures (Bertram *et al.*, 2001; Blumenfeld *et al.*, 2007), and more importantly, facilitate seizure generalization (Bertram *et al.*, 2008). Increasing GABAergic inhibition to the medial-dorsal nuclei can suppress the response to stimulation-induced limbic seizures and further attenuate seizure propagation (Sloan *et al.*, 2011). Taken together, our finding of a greater integrating capacity in the ipsilateral medial-dorsal nuclei of the patients with positive FBTCS history is in line with these concepts. While causality cannot be implicated with cross-sectional data, it is plausible that they are either a maladaptation to FBTCS, or are necessary for to facilitate seizure spread that leads to FBTCS. Note that similar abnormalities were found in both the *current*- and the *remote*-FBTCS group, suggesting that for the latter group, although their FBTCS were under control, they may still possess a vulnerability to FBTCS compared with patients who lack a history of FBTCS. Such shared functional abnormalities may in turn imply additional limbic neurophysiological alterations in these two groups. For instance, although we did not find any direct proof linking MRI-evidenced temporal pathology difference to the PC difference observed here (*Supplementary Result 6*), the *current*- and the *remote*-FBTCS groups indeed included more cases of MRI-negative patients, who could still present with subtle limbic pathologies (Carne *et al.*, 2004).

In addition to an altered thalamocortical network, our findings pointed to a reorganization in the BG–thalamus network specifically in patients with uncontrolled FBTCS, manifested as enhanced interactions between the putamen and the GPi, as well as reduced interactions between the GPi and the thalamus. In animal models, exogenous activation of the striatum can provide a protective effect against seizures (Cavalheiro *et al.*, 1987; Turski *et al.*, 1988; Sabatino *et al.*, 1989; Deransart *et al.*, 1998). In patients, endogenous hypersynchrony between the cortex and the striatum has been observed during seizure termination (Aupy *et al.*, 2019). It is proposed that the striatum modulates the BG output structures, such as the GPi and SN *pars reticulata*, thereby restoring thalamocortical synchrony and terminating the seizure (Aupy *et al.*, 2019). Although exactly *when* the striatum becomes involved remains controversial (Aupy *et al.*, 2019), some scholars suggest the striatal response may be specifically associated with seizure spread and secondary generalization (Rektor *et al.*, 2002, 2011; Popovic *et al.*, 2012; Uchida *et al.*, 2013). Conceivably, this process in patients with uncontrolled FBTCS may be triggered more frequently, yielding a maladaptive increase in striatum–GPi integration.

Importantly, the striatum interacts with the GPi through both direct and indirect pathways, although their distinct contributions to the proposed anticonvulsive process remain unclear (Vuong and Devergnas, 2018). Here, through a series of “disconnection” simulations, we are able to articulate the importance of the direct pathway over the indirect pathway (Figure 5) in revealing FBTCS related changes in interregional integration. In the direct pathway, the striatum projects inhibitory GABAergic efferent connections directly to the GPi, which in turn provides tonic inhibition to the thalamus through GABAergic projections into the lateral and ventral thalamic nuclei (Lanciego *et al.*, 2012). Accordingly, one of the inherent products of an elevated putamen– GPi interaction would be GPi inhibition and, consequently, the reduction of its interaction with downstream targets, such as the lateral-ventral nuclei, matching the anti-correlations we observed between the two sets of integration estimates. In addition, GPi–posterior nuclei integration was also reduced in the *current*-FBTCS patients. Through our “disconnection” simulations, we demonstrated that this difference largely depended upon functional connections from the GPi to the lateral-ventral nuclei, and from the latter to the posterior nuclei, but not upon the direct GPi— posterior nuclei connections (Figure 6). Notably, these findings are tightly aligned with the physiological connections among these structures: the ventrolateral GPi receives input from the putamen and projects to the lateral-ventral thalamic nuclei, while the posterior thalamic nuclei do not receive direct projections from GPi (Alexander *et al.*, 1986). Furthermore, “disconnecting” the direct pathway removed group differences not only in the putamen–GPi integration, but also in both of the GPi–thalamus integrations. Accordingly, we suspect that these specific integration changes in the *current*-FBTCS patients may reflect one unified maladaptive reorganization process. Such a reorganization pattern was only observed in patients with uncontrolled FBTCS, potentially associated with the greater disease burden in this group (*i.e.*, extra secondary seizure generalization). Alternatively, it may also imply that such reorganization is reversible, and can be restored once the FBTCS is under control.

Decreases in the GPi–thalamus interactions imply that in the *current*-FBTCS patients, the tonic thalamic inhibition from the GPi may be weakened. To our knowledge, there exists no direct evidence linking FBTCS with chronic disinhibition of the thalamus. Through single-unit microelectrode recordings from anterior thalamic neurons, Hodaie *et al.* (2006) observed an atypical “hyperactive” firing mode in awake patients with a history of generalized tonic-clonic seizures, which is consistent with our theory. In addition, FBTCS has been associated with greater reduction in the structural integrity of the thalamus (Yang *et al.*, 2017) and thalamocortical connections (He *et al.*, 2015; Keller *et al.*, 2015; Chen *et al.*, 2019), findings which may also be interpreted as the consequences of tonic pathological thalamic activity.

Among the anatomical thalamic nuclei included in our lateral-ventral and posterior ROIs, the centromedian nuclei and medial pulvinar may be particularly relevant to FBTCS control in TLE, but probably through different roles. The medial pulvinar has reciprocal connections with both the medial and lateral part of the temporal lobe (Mauguiere and Baleydier, 1978; Baleydier and Mauguiere, 1985; Insausti *et al.*, 1987; Rosenberg *et al.*, 2009). During temporal lobe seizures, ictal involvement of the medial pulvinar has been consistently recorded (Guye *et al.*, 2006; Rosenberg *et al.*, 2006). High-frequency stimulation (*i.e.*, inhibition) of the medial pulvinar can reduce the severity of hippocampal seizures and shorten its generalization (Filipescu *et al.*, 2019). Therefore, a theoretical reduction in its tonic inhibition could lead to a higher susceptibility to *seizure propagation* from the temporal lobe (Rosenberg *et al.*, 2009). The centromedian nuclei is also considered a prominent target for brain stimulation in epilepsy, though more for its role in gatekeeping and rhythm-generating activities (Lega *et al.*, 2010). High-frequency stimulation of the centromedian nuclei was found mostly efficient in controlling primary and secondary generalized seizures (Velasco *et al.*, 1987, 2006; Fisher *et al.*, 1992). Accordingly, the theoretical reduction in its tonic inhibition could lead to a higher susceptibility to *seizure generalization* (Velasco *et al.*, 1989). Our findings imply that BG stimulation, such as GPi stimulation (Fisher and Velasco, 2014), may restore the tonic inhibition for all the downstream thalamic nuclei (Aupy *et al.*, 2019), leading to antiepileptic effects in TLE patients with uncontrolled FBTCS.

Several methodological considerations are also pertinent to this study. First, due to the limited spatial resolution of rsfMRI, we did not further parcellate the SN into *pars reticulata* and *pars compacta*, acknowledging their distinct rules in the BG pathways. Second, the reliability of our findings depends heavily upon quality control during data processing. Therefore, we adopted the strategy proposed in Satterthwaite *et al.* (2013) using a combination of a sophisticated nuisance regression model and scrubbing. This strategy is sufficient for head motion control (Ciric *et al.*, 2017), even in studies based on short rsfMRI scans (Gu *et al.*, 2015; Xia *et al.*, 2018). However, it is also associated with lower degrees of freedom, which can harm statistical power during FC estimation (Parkes *et al.*, 2018). We also used linear regression to mitigate the effects of confounding factors including demographical, clinical, data quality, and brain structural variations among these patients. Nevertheless, we cannot fully rule out the existence of residual nonlinear influence from these confounders, *e.g.*, the underlying etiology of epilepsy as expressed by differences in pathology among groups. Third, while our “disconnection” simulation conveniently permits the comparison and contrast of different possible pathways, caution should be taken when interpreting these results, as FC essentially measures cross-regional couplings that can be attributed to both direct and indirect physiological connections. Accordingly, instead of mimicking an actual physical disconnection of specific anatomical tracts, this analysis probes putative functional interactions as a blueprint for brain stimulation. Fourth, our conceptual model omitted the connections between the BG and the prefrontal cortex, which is upstream of the BG pathways (Alexander *et al.*, 1986, 1990; Smith *et al.*, 1998). Whilst we made some efforts to test these connections (*Supplementary Result 7*), we did not find FBTCS specific differences. Fifth, due to the lack of simultaneous EEG, we cannot completely rule out the existence of interictal activity during our scan, given its well-known and common effects in TLE (Laufs *et al.*, 2007). Since functional connections including thalamic and BG regions have been shown to be very sensitive to interictal activity (Ibrahim *et al.*, 2014; Shamshiri *et al.*, 2017), the relationship between interictal activity and the functional organization of these networks warrants further investigation. Finally, we should note that some antiepileptic drugs (AED) can influence the concentration of certain neurotransmitters such as GABA, hence potentially impacting the BG–thalamus interaction (Caciagli *et al.*, 2017). Unfortunately, AED regimen heterogeneity (type, dosage, number of AEDs) prevented further testing of these effects. Given the mixed distribution of AEDs across the patient groups (*Supplementary Table S5*), it seems unlikely that the observed difference could be traced to a specific medication.

In summary, we provide evidence that FBTCS imprints on the BG—thalamocortical loops in TLE patients. We demonstrate how a positive history of FBTCS relates to promoted thalamus mediated cortico-cortical communication, and how uncontrolled FBTCS relates to reduced BG— thalamus interactions through the direct pathway. While future longitudinal studies may elaborate more on the underlying causality, our findings have brought additional clarity regarding specific links in the neuronal scaffolding underlying the presence of FBTCS. These mechanistic underpinnings may guide the development of new treatment strategies which may be used to either enhance or diminish interactions in specific circuits.

## Acknowledgement

The authors thank Drs. Gaelle Doucet and Dorian Pustina for help in data acquisition. We thank Drs. Arian Ashourvan, and Maxwell Bertolero for their suggestions. The authors thank all the healthy controls and patients with epilepsy, kept anonymous, who provided data for this study.

## Abbreviations

AED: antiepileptic drugs
BG: basal ganglia
FBTCS: focal to bilateral tonic-clonic seizures
FC: functional connectivity
FIAS: focal impaired awareness seizures
GPe: globus pallidus externus
GPi: globus pallidus internus
HCP: Human Connectome Project
ICA: independent component analysis
PC: participation coefficient
ROI: region of interest
rsfMRI: resting-state functional MRI
SN: substantia nigra
STN: subthalamic nucleus
TLE: temporal lobe epilepsy

