## Supplementary for "Disrupted basal ganglia—thalamocortical loops in focal to bilateral tonic-clonic seizures"

### Supplementary Methods

#### *Imaging Acquisition and Preprocessing*

*Human Connectome Project Data*

The independent resting-state functional MRI (rsfMRI) dataset used for the masked independent component analysis (ICA) based functional parcellation was collected as part of the Washington University-Minnesota Consortium Human Connectome Project (HCP) (https://www.humanconnectome.org/) (Van Essen *et al.*, 2013). All participants gave informed consent consistent with policies approved by the Washington University Institutional Review Board. Whole-brain echo-planar imaging acquisitions were acquired with a 32-channel head coil on a modified 3T Siemens Skyra Scanner (72 slices; TR = 720 ms, TE = 33.1 ms, flip angle = 52°, FOV = 208 × 180 mm, 2.0 mm isotropic voxels, with a multi-band acceleration factor of 8) (Uğurbil *et al.*, 2013). Two runs of a 15-minute resting-state acquisition protocol were used for each participant [see details in (Smith *et al.*, 2013)].

Our analyses were based on the ICA-FIX rsfMRI data provided by the HCP. The "100 Unrelated Subjects" (N=100) subset from the "1200 Subjects" HCP release was used (54 females, age range: 22 to 36 years). In brief, these data were preprocessed with a minimal preprocessing pipeline (Glasser *et al.*, 2013), including motion and distortion correction, normalization to Montreal Neurological Institute (MNI) standard space, and minimal high-pass filtering (2000 seconds). Nuisance regression was performed using FMRIB's ICA based X-noisifier (FIX) (Salimi-Khorshidi *et al.*, 2014; Griffanti *et al.*, 2017), which regresses out independent components classified as noise and 24 motion parameter timeseries estimated through motion correction procedures. The data were subsequently bandpass filtered from 0.01 to 0.08 Hz using 3dBandpass from Analysis of Functional NeuroImages (AFNI, https://afni.nimh.nih.gov).

*Onsite Imaging Data*

All participants were scanned on a 3-T X-series Philips Achieva clinical MRI scanner (Amsterdam, the Netherlands) at Thomas Jefferson University Hospital. A total of five minutes of a resting-state scan was collected from all participants with a single shot echoplanar gradient echo imaging sequence acquiring T2* signals (120 volumes; 34 axial slices acquired parallel to the anterior, posterior commissure line; TR = 2.5 s, TE = 35 ms, flip angle = 90°, FOV = 256 × 256 mm, 128 × 128 data matrix voxels, in-plane resolution = 2 mm × 2 mm, slice thickness = 4 mm). Participants lay in a foam pad to comfortably stabilize the head, were instructed to remain still throughout the scan (i.e., not fall asleep), and were asked to keep their eyes closed during the entirety of the scan. Each imaging series started with three discarded scans to allow for signal stabilization. Prior to collection of the T2* images, T1-weighted images (180 slices) were collected using an MPRage sequence (256 × 256 isotropic 1mm voxels; TR = 640 ms, TE = 3.2 ms, flip angle = 8°, FOV = 256 × 256 mm) in positions identical to the functional scans to provide an anatomical reference. The in-plane resolution for each T1 slice was 1 $mm^3$ (axial oblique).

These imaging data were preprocessed using a combination of neuroimaging tools including Statistical Parametric Mapping 12 (SPM 12, http://www.fil.ion.ucl.ac.uk/spm/software/spm12), AFNI, FMRIB Software Library (FSL v5.0.11, https://fsl.fmrib.ox.ac.uk/fsl/fslwiki/), and FreeSurfer v6.0 (https://surfer.nmr.mgh.harvard.edu). Briefly, as recommended in Power *et al.* (2017), we started our preprocessing with head motion parameters estimation, followed by slice timing correction to adjust for variable acquisition time over slices in a volume. Next, a six-parameter variance cost function rigid body affine registration was used to realign all images within a session to the first volume. Individual structural images (T1-weighted MPRage) were co-registered to the mean functional image using a rigid-body transformation. The transformed structural images were then segmented into gray matter, white matter, and CSF (Ashburner and Friston, 2005). A nuisance regression were performed to regress out the head motion parameters, white matter, CSF and global signals, with all of their derivatives using a 36-parameter model (Satterthwaite *et al.*, 2013), as well as the linear and quadratic trends of the data. Additionally in this model, we also added spike regressors to further reduce the influence of head micromovement. Following (Satterthwaite *et al.*, 2013), we defined spikes as volumes with a framewise displacement (FD) (Jenkinson, 2002) value over 0.25, and generated a separate nuisance regressor for each spike. Accordingly, the number of spikes and corresponding regressors varied over participants. To further ensure the data quality, participants either with excessive head motion (3 mm or degree in max head motion) or with more than 10% of data identified as spikes were excluded and not considered in the final count. This choice ensured that the uncontaminated data in all remaining participants spanned no less than 4 minutes, as recommended by previous studies (Van Dijk *et al.*, 2012; Satterthwaite *et al.*, 2013).

Functional images were then normalized into MNI space in two steps. First, functional images were linearly registered to the structural image using boundary-based registration (BBR) (Greve and Fischl, 2009). We then performed both a linear (Jenkinson and Smith, 2001) and a nonlinear

(Andersson *et al.*, 2007) transformation to register the structural image to the MNI space, so that the resultant warp parameters can be applied to the functional images. The normalized functional images were spatially resampled to a 2 mm voxel resolution, and subsequently bandpass filtered from 0.01 to 0.08 Hz. Because the basal ganglia (BG) and thalamus are relatively small structures, we refrained from global spatial smoothing in order to avoid signal blurring. Lastly, the volumes identified as spikes were scrubbed and excluded from the subsequent functional connectivity analyses.

#### *Distribution Pattern of Thalamocortical Connections*

To test our hypothesis that the thalamus plays a role as a connector hub in the brain to facilitate secondary generalization of seizures, we estimated the participation coefficient (PC) of each thalamic parcel based on its thalamocortical connections (Hwang *et al.*, 2017). Specifically, we were interested in the intrinsic functional connectivity (FC) between thalamus and the cerebrum, and how uniformly these thalamocortical connections were distributed across known cortical networks. Briefly, we extracted the mean time-series from 200 cortical ROIs functionally defined by Schaefer *et al.* (2018). The 200 ROIs were grouped into 7 cortical resting-state networks (RSNs) according to Yeo *et al.* (2011), so that each thalamocortical connection could be related to one of the RSNs. In line with the predominantly unilateral organization of thalamocortical connections (Jones, 2007), we estimated the thalamocortical FC unilaterally between each thalamic parcel and cortical ROI using a Pearson correlation coefficient, yielding a 7 (thalamic parcels) by 100 (cortical ROIs) matrix for each hemisphere. We further removed the unreliable connections (i.e., correlations with an FDR-corrected $P$ value > 0.05) by setting their weights to zero, and took the absolute value of all remaining connection weights. We defined the PC of region $i$ as

$$PC_i = 1 - \sum_{S=1}^{N_M} \left(\frac{k_{is}}{k_i}\right)^2 , \quad (1)$$

where $K_i$ is the sum of the connectivity weight of region $i$, $K_{is}$ is the sum of the connectivity weight between region $i$ and the cortical network $s$, and $N_M$ is the total number of networks. Accordingly, a region with more uniformly distributed connections will present a PC closer to 1, and in contrast, a region with more varyingly distributed connections will present a PC closer to 0. The PC value of each thalamic parcel was estimated and further scaled to [0,1] by dividing the theoretical maximum calculated as $PC_{max} = 1 - 7 \times \left(1/7\right)^2 = 0.857$ (a perfect uniform distribution across 7 RSNs). As a measure independent of the node's degree, PC is less biased by the number of ROIs within each cortical network (Power *et al.*, 2013). Furthermore, we validated our PC analysis using matrices involving binary matrices or bilateral thalamocortical connections (7×200) and found similar results (Supplementary Results 2).

#### *Community Detection based Interregional Integration*

To investigate the interaction between the BG and the thalamus, we constructed the absolute Pearson correlation matrix between the BG and the thalamus ROIs for each hemisphere, respectively, keeping in line with the unilateral nature of the BG–thalamus circuits (Alexander *et al.*, 1986; Parent and Hazrati, 1995). We further applied a similar strategy to remove statistically unreliable connections (i.e., correlations with an FDR-corrected $P$ value > 0.05) from each matrix. Accordingly, each matrix represented the unilateral BG–thalamus network, with each ROI represented as a node and pairwise correlations between ROI time-series represented as weighted edges.

From these networks, we sought to parameterize the intrinsic functional interactions among these regions using tools from network science, namely, community detection (Fortunato, 2010). Communities correspond to groups of nodes that are more strongly connected to one another than would be expected by chance alone (Fortunato, 2010). Accordingly, nodes assigned to the same community will tend to interact with each other more strongly than with nodes from other communities. To detect communities from these networks, we used a generalized Louvain-like method (Blondel *et al.*, 2008) to optimize a modularity quality function (Newman and Girvan, 2004; Newman, 2006; Reichardt and Bornholdt, 2006):

$$Q = \frac{1}{2m} \sum_{ij} \left(A_{ij} - \gamma V_{ij}\right) \delta\left(g_i, g_j\right) , \tag{2}$$

where for the edge between node $i$ and node $j$, $A_{ij}$ is the observed weight, and $V_{ij}$ is the expected weight under the Newman-Girvan null model (Newman and Girvan, 2004), given by (2):

$$V_{ij} = \frac{k_i k_j}{2m} , \tag{3}$$

where $m = \frac{1}{2} \sum_{ij} A_{ij}$ is the total edge weight in the network, and $k_i$ and $k_j$ refer to the strength of node $i$ and node $j$, respectively. The quantities $g_i$ and $g_j$ represent the community assignments of node $i$ and node $j$, respectively, and the Kronecker delta function $\delta\left(g_i, g_j\right) = 1$ if $g_i = g_j$ and 0 otherwise. The parameter $\gamma$ is a structural resolution parameter (Reichardt and Bornholdt, 2006). As a standard and in absence of an *a priori* hypothesis, we set $\gamma$ to 1. However, to demonstrate that our results are robust to different choices for the value of this resolution parameter, we reanalyzed our data with different $\gamma$ values ($\gamma = [0.95, 1.05]$), obtaining qualitatively similar results (*Supplementary Result 3*).

Due to the stochastic nature of the algorithm and near degeneracy of the underlying modularity landscape (Good *et al.*, 2010), each independent run of the community detection algorithm may provide slightly different partitions of network nodes into communities. Therefore, we performed 1000 optimizations of the modularity quality function for each participant as done in prior work (Bassett *et al.*, 2013; He *et al.*, 2018). All subsequent diagnostics based on the outputs of community detection were estimated at each optimization, and then averaged over all optimizations to generate representative values for each of them.

Once every node from every network was assigned into one community, we used the *module allegiance* matrix *P* to provide a summary of the co-assignment of each pair of nodes into the same community following prior literature (Bassett *et al.*, 2015; Mattar *et al.*, 2015). Specifically for each network during each optimization, each element $P_{ij}$ of the module allegiance matrix *P* equals 1 if node $i$ and node $j$ are assigned to the same community, and equals 0 if the two nodes are assigned to different communities. Accordingly, the module allegiance matrix represents the preference of intrinsic functional interaction at the pairwise nodal level, which can emerge both among subdivisions from the same anatomical structure, or between components from different anatomical origins. As we were particularly interested in the interactions between the BG and the thalamus, we subsequently grouped the ROIs based on their anatomical origins and let $C = \{C_1, \cdots, C_k\}$ be the anatomical partitions of the network. Then following (Bassett *et al.*, 2015), we defined:

$$I_{k_1,k_2} = \frac{\sum_{i \in C_{k_1}, j \in C_{k_2}} P_{ij}}{|C_{k_1}||C_{k_2}|}, \tag{4}$$

as the *interregional integration* between partition $C_{k_1}$ and partition $C_{k_2}$ when $k_1 \neq k_2$, where $|C_k|$ is the number of nodes in partition $C_k$. The integration values were estimated for each pair of anatomical partitions and averaged over the 1000 optimizations of the modularity quality function. Intuitively, a higher value represented a higher probability of the members from one partition been assigned to the same community with the members from another partition, potentially suggesting a higher functional interaction between these two anatomical structures.

#### *Random Network Null Model*

To further validate that the results from group-level comparisons were due to the differences in participants' topological organization of the BG–thalamus network, we employed a random network null model as a benchmark (Rubinov and Sporns, 2011). This procedure began with a real network and then iteratively rewired and reassigned each weighted edge at random, preserving both the degree and strength distribution of the true network. The process was performed multiple times for each network, such that each edge was 'rewired' approximately 20 times (Maslov and Sneppen, 2002). In total 100 random network null models were generated for each real network and underwent the same community detection procedure (again with 1000 optimizations), yielding a set of separate interregional integration measures. Identical group-level comparisons were made on these null model integration values.

#### *Permutation-based Statistical Testing*

To minimize the bias of data distribution to our statistical inferences and correct for multiple comparisons when appropriate, we implemented a permutation-based method as our main statistical strategy throughout this study (Groppe *et al.*, 2011). Individual permutation-based statistical testing permits the inference of the probability of the observed statistic (e.g., $t$ value), from a distribution of the same statistic estimated from massive instances of the same samples with their group identities permuted (Good, 2005). In many cases we wish to apply a permutation-based test to scenarios with multiple comparisons, i.e., comparing multiple within-subject variables across the same groups of subjects. In this case, we can expand the traditional approach by applying a "$t_{max}$" principle to adjust the estimated $P$-values of each variable for multiple comparisons by controlling the family-wise error rate (Blair and Karniski, 1993). Briefly, the observed statistic for each variable is compared to the distribution of the most extreme statistic across the entire family of tests for each possible permutation. This procedure corrects for multiple comparisons because the distribution of the most extreme statistics automatically adjusts to reflect the increased chance of false discoveries due to an increased number of comparisons (Groppe *et al.*, 2011). We performed 1,000,000 permutations each time we implemented this strategy to ensure high precision during $P$-value estimation, and we used either $t$ or $F$ statistics when appropriate.

### Supplementary Results

#### *Data Quality Comparisons*

Before testing our main hypotheses based on intrinsic FC, we examined two main factors that may also bias the subsequent analyses, namely the anatomical structure where fMRI signals were extracted and the quality of the data regarding head motion control and community detection. We first tested whether the patient groups differed anatomically in these subcortical regions. Through FreeSurfer, volumetric statistics for 3 striatum subdivisions (caudate, putamen, and ventral striatum), GP, thalamus, hippocampus and amygdala were derived from each hemisphere, as well as a summary measure of total subcortical gray matter volumes. With a permutation-based $F$ test, we found no significant group differences in these subcortical volumes (*Supplementary Table S4*), after regressing out age, sex, handedness, seizure focus side, and estimated intracranial volume. To minimize the effect of the volumetric variances, the total subcortical gray matter volume was still added as a confounding factor, although it was unlikely that any subsequent group-level differences could be mainly attributed to the volumetric differences in these subcortical structures.

Next, we tested whether there was any difference in data quality across the three groups. As shown in *Supplementary Table S2 and S3*, both the head micromovement, measured as mean FD, and the number of spikes detected and scrubbed from the data were balanced across all experimental groups. Furthermore, the density of the FC matrix after FDR correction was also consistent across groups and hemispheres. Moreover, the community detection yielded similar performances across groups, evidenced by comparable values of the modularity index Q and maximum number of communities. Despite the lack of any group differences, we added the mean FD, number of spikes, and network density as confounding factors to further reduce their potential influence on our findings. Note that, modularity index Q and maximum number of communities were not included because as parallel products, it is impossible for them to pose any causal influence on the network properties.

#### *Reproducibility Tests on Distribution Patterns of Thalamocortical Connections*

To test for the robustness of our findings in distribution patterns of thalamocortical connections against the selection of analytical strategies, we performed three additional reproducibility analyses. First, we sought to determine whether the group difference in thalamic participant coefficient (PC) could still be found without regressing out confound variables. We therefore compared the raw PC values directly estimated from the FDR-corrected, absolute thalamocortical functional connectivity (FC) matrices across the three patient groups, using the same permutation-based $F$ test. We found a significant group difference only in the ipsilateral medial-dorsal nuclei ($F(2,93)=10.392$, $P_{corr}=4.5\times10^{-4}$; others: $F(2,93)<1.958$, $P_{corr}>0.705$) similar to the result that we reported in the main text. Second, we tested whether the way we estimated the PC using the weighted FC matrix would bias our observations. So instead, here we re-estimated the PC after binarizing the FC matrix, regressed out the confound variables, and compared across the patient groups. Again, we observed a similar group difference only in the ipsilateral medial-dorsal nuclei ($F(2,93)=6.553$, $P_{corr}=0.014$; others: $F(2,93)<2.363$, $P_{corr}>0.558$). Third, we tested whether the result held if the bilateral functional thalamocortical connections were taken into account. While direct cross-hemisphere thalamocortical connections are physiologically impossible, functional synchrony, as measured by FC, can be plausible via indirect physiological connections. Specifically, the thalamocortical FC matrix was generated bilaterally (7×200) instead of unilaterally (7×100) for each thalamus. From the FDR-corrected, absolute-value transformed new matrix, we estimated and compared the thalamic PC after confound regression. As expected, we still observed the group difference only in the ipsilateral medial-dorsal nuclei ($F(2,93)=6.188$, $P_{corr}=0.022$; others: $F(2,93)<2.003$, $P_{corr}>0.683$). Taken together, these reproducibility tests indicate that the reported group differences in the distribution patterns of thalamocortical connections are robust against the choices of analytical strategies such as confound regression, weight of the FC matrix, and the inclusion of contralateral thalamocortical FCs.

#### ***Reproducibility Tests on Integration between Basal Ganglia and Thalamus***

To test for the robustness of our findings in integration between basal ganglia and thalamus against the selection of analytical strategies, we performed two additional reproducibility analyses. First, we sought to determine whether the group differences of integration in the BG—thalamus network could still be found without regressing out confound variables. We therefore compared the raw integration values directly estimated from the FDR-corrected, absolute BG—thalamus FC matrices across the three patient groups, using the same permutation-based $F$ test. We found significant group differences in the integration of ipsilateral striatum–GP pair ($F(2,93)=6.666$, $P_{corr}=0.039$) and ipsilateral GP–thalamus pair ($F(2,93)=8.905$, $P_{corr}=0.006$; others: $F(2,93)<3.033$, $P_{corr}>0.630$), similar to the results that we reported in the main text. Second, we introduced small variances in the resolution parameter $\gamma$ used for community detection, and we explored whether the aforementioned integration differences would remain. Since we used a setting of $\gamma = 1$ in the main text, here we tested both a smaller ($\gamma = 0.95$) and a larger ($\gamma = 1.05$) setting. Then we re-estimated the integration, regressed out confound variables, and compared across the patient groups. After setting γ to 0.95, we observed similar results of significant group differences in the integration of the ipsilateral striatum–GP pair ($F(2,93)=7.411$, $P_{corr}=0.021$) and the ipsilateral GP–thalamus pair ($F(2,93)=9.027$, $P_{corr}=0.005$; others: $F(2,93)<4.398$, $P_{corr}>0.246$). Similarly, after setting γ to 1.05, we were able to observe comparable results of significant group differences in the integration of the ipsilateral striatum–GP pair ($F(2,93)=9.100$, $P_{corr}=0.005$) and the ipsilateral GP–thalamus pair ($F(2,93)=8.356$, $P_{corr}=0.009$; others: $F(2,93)<2.461$, $P_{corr}>0.815$). Taken together, these reproducibility tests indicate that the reported group differences in the integration between basal ganglia and thalamus are robust to the choices of analytical strategies such as confound regression and selection of the resolution parameter during community detection.

#### *Relationship between Integration, Pairwise FC Intensity and FBTCS History*

The absence of group difference in the pairwise FC intensity suggested that the collinearity between the pairwise FC intensity and our grouping scheme (i.e., FBTCS history difference) was generally low, hence further allowing us to explore to what extent the observed interregional integration differences can be explained by the pairwise FC intensity and the FBTCS history difference, respectively. Accordingly, we formulated a linear regression model with the integration value as dependent variable, the 20 pairwise FC intensity estimates as independent variables entered in a stepwise fashion, and the group index "force entered" at the last step. When taking the striatum–GP integration as the dependent variable, we found a significant model with three independent variables included ($R^2=0.403$, $F(3,92)=20.714$, $P=2.4\times10^{-10}$): the ipsilateral striatum–GP ($\beta=0.554$, $t=6.231$, $P=1.4\times10^{-8}$, VIF=1.219) and GP–thalamus ($\beta=-0.405$, $t=-4.646$, $P=1.1\times10^{-5}$, VIF=1.169) FC intensity, plus the group index ($\beta=0.217$, $t=2.623$, $P=0.010$, VIF=1.053). When taking the GP–thalamus integration as the dependent variable, we also found a significant model with three independent variables included ($R^2=0.460$, $F(3,92)=26.102$, $P=2.6\times10^{-12}$): the ipsilateral striatum–GP ($\beta=-0.472$, $t=-5.581$, $P=2.4\times10^{-7}$, VIF=1.219) and GP–thalamus ($\beta=0.523$, $t=6.307$, $P=9.8\times10^{-9}$, VIF=1.169) FC intensity, plus the group index ($\beta=-0.293$, $t=-3.725$, $P=3.4\times10^{-4}$,VIF=1.053). In both models, there was significant remaining variance explained by the FBTCS history difference, after the most relevant pairwise FC intensities were taken into account.

#### *Reproducibility Test on Globus Pallidus Centered Integration*

To assess the robustness of our findings in integration among striatum, GP and thalamus against the utility of confound regression, we tested whether the group differences of integration reported in the main text could be reproduced without regressing out the confound variables. Specifically, we compared the raw integration values across the three patient groups with the same permutation-based *F* test. We observed the same overall effects, including no significant group differences among the pairwise integrations with GPe ($F$'s(2,93)<3.685, $P_{corr}$'s>0.371), and significant group differences in the integration of the ipsilateral putamen–GPi pair ($F$(2,93)=9.025, $P_{corr}$=0.006), the ipsilateral GPi– lateral-ventral nuclei pair ($F$(2,93)=6.622, $P_{corr}$=0.037), and the ipsilateral GPi–posterior nuclei pair ($F$(2,93)=8.247, $P_{corr}$=0.010; others: $F$'s(2,93)<3.316, $P_{corr}$'s>0.484). These results indicate that the reported group differences in the GPi integration are robust against the utility of our confound regression strategy.

#### *Relationship between MRI-Evidenced Pathology and thalamic PC*

Compared to the *none*-FBTCS group, both the *remote*- and *current*-FBTCS groups presented a higher proportion of MRI-negative cases, implying potential differences in subtle limbic pathologies (Carne *et al.*, 2004). They also both presented elevated PC values at the ipsilateral medial-dorsal (MD) thalamic nuclear group, which is heavily implicated in limbic circuitry (Alexander *et al.*, 1986, 1990; Dolleman-Van der Weel *et al.*, 1997; Bertram and Zhang, 1999; Behrens *et al.*, 2003). Here, we specifically explored such co-existence of group differences, to determine whether the PC difference was driven by the etiology/pathology differences other than the FBTCS history.

First, we performed a two-way ANOVA, using the PC values from the ipsilateral MD as a dependent variable, with both FBTCS group and MRI-evidenced temporal pathology (to be henceforth referred to as "MRI-pathology" for short) as fixed factors. We followed the same steps and procedure outlined in the main text, excepting for the exclusion of the MRI-pathology from the confounder list since it is now a variable of interest. In line with our main results, we found a significant main effect of FBTCS group [$F_{(2,87)}$=6.516, $P$=0.002]. However, the main effect of MRI-pathology was not significant [$F_{(2,87)}$=1.180, $P$=0.312], nor was the interaction between MRI-pathology and FBTCS group [$F_{(4,87)}$=0.629, $P$=0.643]. Therefore, it is unlikely that MRI-pathology is the main factor driving the PC difference at the ipsilateral MD, either alone or through its interaction with FBTCS history.

Second, since the true pathological nature of MRI-negative cases was unclear, we decided to focus on a patient subgroup with well-characterized limbic pathology, *i.e.*, TLE-HS, and we evaluated whether the previously-detailed PC difference could be revealed in this homogeneous subgroup. Of note, the proportion of HS-positive cases was balanced across the three patient groups (*none-*/*remote-*/*current-*=14/16/11, $\chi^2$=1.618, $P$=0.445), justifying a valid subgroup analysis. We followed the same settings as in the main text, except for the exclusion of the MRI-evidenced pathology from the confounder list as we were only testing HS-positive patients here. After correction for multiple comparisons (*FWE*), we found a significant group difference for PC of the ipsilateral MD only, with the same permutation test [$F_{(2,38)}$=6.762, $P_{corr}$=0.020 ($P_{unc}$=0.003)]. This result indicates that even in a homogenous subgroup of patients with the same limbic pathology, such PC difference can still be detected, and can ultimately be attributed to propensity for secondary generalization.

Collectively, these additional analyses do not suggest that MRI-evidenced temporal pathology may significantly affect PC differences at ipsilateral MD in our data.

#### *BG-prefrontal connections in patients with different FBTCS history*

As the upstream of the BG pathways (Alexander *et al.*, 1986, 1990; Smith *et al.*, 1998), the connections between BG and prefrontal cortex may also be affected by the prevalence of FBTCS. It is intriguing to "close the loop" and test the interaction between BG, thalamus, and the prefrontal areas known to be connected to BG, such as premotor cortex, supplementary motor area (SMA), frontal-eye-field (FEF), and anterior cingulate cortex (ACC) (Alexander *et al.*, 1986, 1990).

We began by locating these prefrontal ROIs using the Brainnetome atlas, a multimodal neuroimaging-based parcellation frequently used for functional MRI investigations, which has optimal spatial granularity and Brodmann area landmarks (Fan *et al.*, 2016). In each hemisphere, this atlas provided five parcels for BA6 (premotor cortex and SMA), one parcel for lateral BA8 (FEF), and four parcels for BA32/24 (ACC). Time series were extracted from these 20 ROIs (10 for each hemisphere) from the preprocessed rsfMRI data as detailed in the main text. Next, for each hemisphere, we added these 10 prefrontal ROIs to the 21 subcortical ROIs, and derived a Pearson correlation matrix for each subject. Following the same pipeline as in the main analysis, we further removed the unreliable connections (*e.g.*, correlations with an FDR-corrected $P$ value > 0.05) by setting their weights to zero, and we took the absolute value of all remaining connection weights for all of the matrices.

With these new data in hand, we evaluated the integration between the subcortical regions and the prefrontal areas in the same way as described in the main text. In brief, we performed community detection on this new matrix, and estimated the integration between the original five partitions (striatum, GP, SN, STN and thalamus) and the prefrontal partition (five pairs per hemisphere, namely, striatum-prefrontal, GP-prefrontal, SN-prefrontal, STN-prefrontal, and thalamus-prefrontal), for each hemisphere and each subject. We then compared these pairwise integration values both ipsilaterally and contralaterally, both with and without the regression of confound variables. We found that none of these pairwise integration values were significantly different across the patient groups with different FBTCS history, either with [$F$'$s_{(2,93)}<1.852$, $P_{unc}$'s>0.163] or without [$F$'$s_{(2,93)}<1.192$, $P_{unc}$'s>0.308] confound regression. Note that all $P$-values reported here were not corrected for multiple comparisons.

Finally, to confirm our findings, we also tested pairwise FC intensity between the subcortical regions and the selected prefrontal areas in the same way as described in the main text. Consistently, we found similar null results; pairwise FC intensity values were not significantly different across the three patient groups, either with [$F$'$s_{(2,93)}<1.814$, $P_{unc}$'s>0.169] or without [$F$'$s_{(2,93)}<2.115$, $P_{unc}$'s>0.126] confound regression. Note that all $P$-values reported here are not corrected for multiple comparisons.

Thus, these results do not support the hypothesis that patients with different FBTCS history would present with specific differences in their BG/thalamus to selective prefrontal connections. One possible explanation is that epilepsy is associated with a widely distributed pathological network (Tavakol *et al.*, 2019), especially compared to Parkinson's disease, in which abnormalities of BG pathways are more specific. Therefore, disruptions of prefrontal connectivity in TLE may be heterogeneous and likely multifactorial, hence may be less likely to be detected statistically. For instance, disruptions in thalamocortical connections have been more consistently reported for temporal lobe locations but not prefrontal cortex (Keller *et al.*, 2014, 2015). Another possible explanation is that disruptions of prefrontal connectivity in TLE may not be specific to FBTCS. Previously, our group found a marginal thalamocortical FC reduction to the entire prefrontal cortex (not as selective as here) in TLE patients compared to healthy controls, with such an effect comparable in patients with and without FBTCS (He *et al.*, 2015).

### Supplementary Tables

**Table S1.** Basal ganglia – thalamus regions of interests.

| Hemisphere | Region | Subdivision | Size (voxels) | X | Y | Z |
|---|---|---|---|---|---|---|
| L | Caudate | AntInf | 194 | 52.6 | 71.9 | 34.5 |
| L | Caudate | AntSup | 130 | 52.3 | 70.7 | 41.2 |
| L | Caudate | Mid | 172 | 49.1 | 68.9 | 37.5 |
| L | Caudate | Post | 260 | 51.9 | 62.7 | 45 |
| L | Ventral Striatum | | 111 | 50.8 | 67.5 | 31.3 |
| L | Putamen | AntInf | 152 | 56 | 66.4 | 32.6 |
| L | Putamen | AntSup | 191 | 56.5 | 67.4 | 37.5 |
| L | Putamen | MidInf | 135 | 59 | 62.7 | 35.2 |
| L | Putamen | MidSup | 150 | 57.8 | 62 | 40.6 |
| L | Putamen | Post | 184 | 59.6 | 56.8 | 37.6 |
| L | Globus Pallidus | External | 147 | 54.7 | 61 | 35.8 |
| L | Globus Pallidus | Internal | 50 | 54.2 | 59.4 | 34.4 |
| L | Substantia Nigra | | 63 | 49.7 | 54.7 | 30.1 |
| L | Subthalamic Nucleus | | 24 | 50.2 | 56.3 | 33 |
| L | Thalamus | Ant | 188 | 48.4 | 59.7 | 39.8 |
| L | Thalamus | MedDor | 160 | 47.1 | 54.4 | 39.3 |
| L | Thalamus | LatDor | 147 | 50.9 | 55 | 42.8 |
| L | Thalamus | LatPost | 113 | 52.9 | 50 | 41.1 |
| L | Thalamus | VenLat | 209 | 50.7 | 54.9 | 37.1 |
| L | Thalamus | MedPul | 141 | 49.5 | 49.1 | 37.9 |
| L | Thalamus | LatPul | 179 | 54.4 | 48.5 | 36.4 |
| R | Caudate | AntInf | 149 | 37.8 | 72.7 | 34.6 |
| R | Caudate | AntSup | 146 | 36.9 | 71.4 | 41.1 |
| R | Caudate | Mid | 174 | 40.3 | 69.6 | 38.2 |
| R | Caudate | Post | 264 | 37.6 | 63.2 | 45.3 |
| R | Ventral Striatum | | 81 | 40.4 | 67.9 | 31.9 |
| R | Putamen | AntInf | 153 | 35.6 | 68.1 | 32.2 |

| | | | | | | |
|---|---|---|---|---|---|---|
| R | Putamen | AntSup | 171 | 33.1 | 68.9 | 36.6 |
| R | Putamen | MidInf | 188 | 31.1 | 63.8 | 34.6 |
| R | Putamen | MidSup | 208 | 31.8 | 63.2 | 40.1 |
| R | Putamen | Post | 168 | 29.9 | 57.4 | 37.7 |
| R | Globus Pallidus | External | 135 | 35 | 61.7 | 36.2 |
| R | Globus Pallidus | Internal | 51 | 35.5 | 59.6 | 34.9 |
| R | Substantia Nigra | | 55 | 39.8 | 55 | 30 |
| R | Subthalamic Nucleus | | 27 | 39.5 | 56.9 | 32.9 |
| R | Thalamus | Ant | 141 | 41.2 | 59.9 | 38.8 |
| R | Thalamus | MedDor | 205 | 41.8 | 54.7 | 37.8 |
| R | Thalamus | LatDor | 167 | 39.5 | 56.7 | 42.4 |
| R | Thalamus | LatPost | 196 | 36.8 | 53 | 40.7 |
| R | Thalamus | VenLat | 109 | 38.2 | 52.5 | 36.1 |
| R | Thalamus | MedPul | 120 | 39.6 | 48.7 | 39.1 |
| R | Thalamus | LatPul | 199 | 34.9 | 48.4 | 36.8 |

Abbreviations & Definitions: Ant, anterior; Mid, middle; Post, posterior; Inf, inferior; Sup, superior; Med, medial; Lat, lateral; Dor, dorsal; Ven, ventral; Pul, pulvinar. Each voxel is 8 $mm^3$ sized. Coordinates were estimated at the center of gravity in MNI space.

**Table S2.** Quality control of the data.

| **TLE Group** (N) | ***none*-FBTCS** (32) | ***remote*-FBTCS** (32) | ***current*-FBTCS** (32) | ***F*** | ***P*** |
|---|---|---|---|---|---|
| Mean FD | 0.08±0.03 | 0.08±0.02 | 0.07±0.02 | 1.057 | 0.352 |
| Spikes Scrubbed | 2.75±2.97 | 1.88±2.14 | 2.00±2.50 | 1.094 | 0.339 |
| Modularity Index Q | | | | | |
| *Ipsilateral Side* | 0.22±0.05 | 0.23±0.06 | 0.21±0.06 | 0.383 | 0.683 |
| *Contralateral Side* | 0.23±0.04 | 0.22±0.05 | 0.23±0.07 | 0.579 | 0.563 |
| Max num. of Communities | | | | | |
| *Ipsilateral Side* | 3.09±0.44 | 3.15±0.53 | 3.32±0.41 | 1.986 | 0.143 |
| *Contralateral Side* | 3.08±0.62 | 3.22±0.51 | 3.25±0.50 | 0.941 | 0.394 |
| Network Density (%) | | | | | |
| *Ipsilateral Side* | 46.80±8.94 | 47.23±8.56 | 46.73±10.07 | 0.028 | 0.972 |
| *Contralateral Side* | 46.64±9.13 | 47.56±8.80 | 44.60±11.94 | 0.727 | 0.486 |

Continuous variables were presented in mean ± standard deviation. Abbreviations & Definitions: TLE, patients with temporal lobe epilepsy; FBTCS, focal to bilateral tonic-clonic seizure; FD, framewise displacement (Jenkinson, 2002), a measure of head motion.

**Table S3.** Comparison between healthy controls and temporal lobe epilepsy patients.

| **Group** (N) | ***none-*** **FBTCS** (32) | ***remote-*** **FBTCS** (32) | ***current-*** **FBTCS** (32) | **Healthy Controls** (32) | $F/\chi^2$ | ***P*** |
|---|---|---|---|---|---|---|
| Age | 41.75±12.77 | 41.66±16.34 | 37.63±11.52 | 36.47±12.59 | 1.322 | 0.270 |
| Gender (Male/Female) | 14/18 | 16/16 | 17/15 | 16/16 | 0.594 | 0.898 |
| Handedness (R/L/A) | 26/6/0 | 28/3/1 | 26/4/2 | 24/8/0 | 1.641 | 0.650 |
| Mean FD | 0.08±0.03 | 0.08±0.02 | 0.07±0.02 | 0.08±0.02 | 0.700 | 0.554 |
| Spikes Scrubbed | 2.75±2.97 | 1.88±2.14 | 2.00±2.50 | 2.41±3.46 | 0.643 | 0.589 |
| Modularity Index Q | | | | | | |
| *Left Side* | 0.23±0.06 | 0.21±0.04 | 0.23±0.06 | 0.23±0.06 | 1.442 | 0.234 |
| *Right Side* | 0.23±0.06 | 0.22±0.05 | 0.22±0.04 | 0.22±0.05 | 0.467 | 0.706 |
| Max num. of Communities | | | | | | |
| *Left Side* | 3.15±0.46 | 3.06±0.42 | 3.35±0.57 | 3.09±0.56 | 2.093 | 0.105 |
| *Right Side* | 3.11±0.53 | 3.23±0.49 | 3.21±0.55 | 3.05±0.69 | 0.691 | 0.559 |
| Network Density (%) | | | | | | |
| *Left Side* | 46.92±8.71 | 46.95±8.59 | 46.09±9.65 | 48.74±10.25 | 0.458 | 0.712 |
| *Right Side* | 46.52±9.35 | 47.84±8.75 | 45.24±12.36 | 49.24±9.17 | 0.947 | 0.420 |

Continuous variables were presented in mean ± standard deviation. Abbreviations & Definitions: FBTCS, focal to bilateral tonic-clonic seizure; FD, framewise displacement (Jenkinson, 2002), a measure of head motion; Note, since side of ictal onset did not apply to healthy controls, here data were presented as left vs. right hemisphere.

**Table S4.** Subcortical volumetric statistics.

| TLE Group (N) | | *none-* FBTCS (32) | *remote-* FBTCS (32) | *current-* FBTCS (32) | $F$ | $P_{corr}$ |
|---|---|---|---|---|---|---|
| Ipsilateral side | Caudate | 3082±339 | 3281±387 | 3263±491 | 2.292 | 0.606 |
| | Putamen | 4540±438 | 4731±460 | 4683±563 | 1.318 | 0.921 |
| | VS | 550±90 | 561±96 | 587±117 | 1.121 | 0.960 |
| | GP | 1899±199 | 1990±196 | 1972±215 | 1.767 | 0.790 |
| | Thalamus | 6982±673 | 7220±666 | 7255±602 | 1.683 | 0.818 |
| | Hippocampus | 3571±595 | 3594±514 | 3731±695 | 0.651 | 0.998 |
| | Amygdala | 1684±325 | 1657±238 | 1777±303 | 1.488 | 0.878 |
| Contralateral side | Caudate | 3326±332 | 3512±366 | 3472±430 | 2.150 | 0.656 |
| | Putamen | 4712±444 | 4972±540 | 4957±507 | 2.753 | 0.457 |
| | VS | 588±75 | 616±80 | 639±94 | 3.007 | 0.386 |
| | GP | 1864±186 | 1921±185 | 1967±202 | 2.320 | 0.597 |
| | Thalamus | 6631±547 | 6909±694 | 6772±633 | 1.569 | 0.854 |
| | Hippocampus | 4096±519 | 4087±482 | 4218±410 | 0.761 | 0.995 |
| | Amygdala | 1886±274 | 1917±227 | 1938±247 | 0.357 | 1.000 |

Volumetric statistics were corrected for age, sex, handedness, seizure focus side, and estimated intracranial volume (mean ± standard deviation). Abbreviations & Definitions: TLE, patients with temporal lobe epilepsy; FBTCS, focal to bilateral tonic-clonic seizure; VS, ventral striatum; GP, globus pallidus.

**Table S5.** Current antiepileptic drugs (AED) by categories and counts.

| | VGNC (+/-) | GABAa (+/-) | SV2a (+/-) | CRMP2 (+/-) | Multi-Action (+/-) | VGCC (+/-) | AED count (1/2/3) |
|---|---|---|---|---|---|---|---|
| *none*-FBTCS | 16/16 | 0/32 | 12/20 | 11/21 | 6/26 | 0/32 | 18/13/1 |
| *remote*-FBTCS | 19/13 | 4/28 | 13/19 | 6/26 | 8/24 | 3/29 | 11/18/3 |
| *current*-FBTCS | 16/16 | 3/29 | 11/21 | 13/19 | 3/29 | 2/30 | 15/15/2 |
| $\chi^2$ | 0.75 | N.A. | 0.27 | 3.78 | 2.72 | N.A. | N.A. |
| *P* | 0.69 | 0.16* | 0.88 | 0.15 | 0.26 | 0.36* | 0.48* |

Category was defined by the presumed mechanism of action of each AED. For simplification, when sorted by category, multiple AEDs from the same category for one patient were considered as one count. When sorted by AED count, multiple AEDs from the same category for one patient were considered separately. Neither the dosage nor the time or times of AED consumption were taken into account. * Fisher's exact test performed instead, since more than 20% of cells had an expected count less than 5. Abbreviations: VGNC, voltage-gated $Na^+$ channel blockage; GABAa, GABAa agonist; SV2a, SV2a receptor mediated; CRMP2, CRMP2 receptor mediated; VGCC, voltage-gated $Ca^{++}$ channel blockage.

### Supplementary Figures

#### Supplementary Figure 1

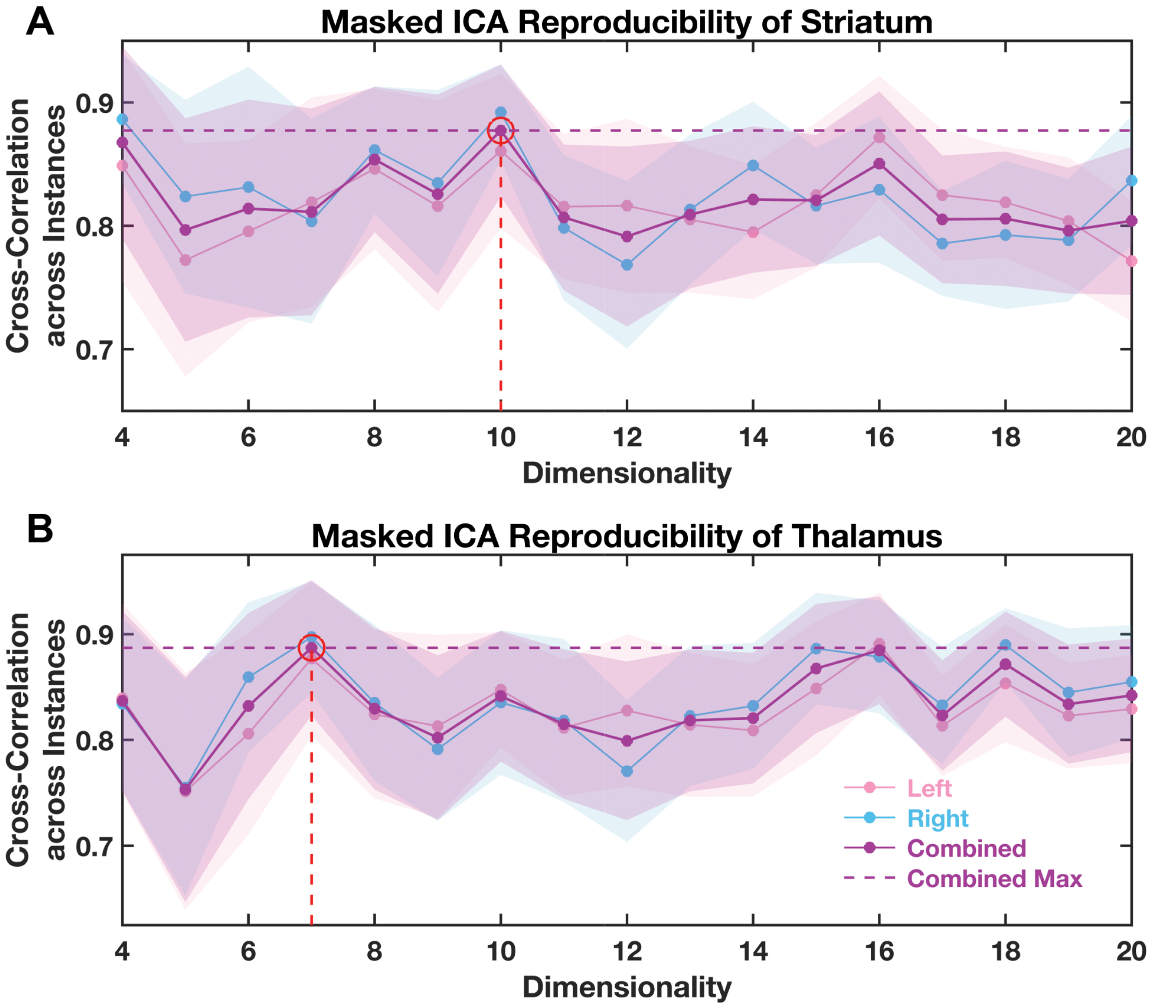

**Supplementary Figure 1. Masked independent component analysis (ICA) reproducibility analyses.** Specifically, the 100 HCP subjects were randomly assigned into halves for 50 instances. For each instance, masked ICA was performed on each half of the samples independently as the number of independent components (ICs) ranged from a dimensionality of 4 to a dimensionality of 20. Spatial cross-correlations were calculated between the ICs extracted from each half of the samples at each setting for the dimensionality (and therefore different number of ICs) and averaged across instances. Higher mean cross-validation correlation suggests higher stability of ICA decompositions. When taking into consideration both hemispheres, the local maximum of the cross-correlation across instances can be found at a dimensionality of 10 for the striatum (panel A) and of 7 for the thalamus (panel B). Left, mean cross-correlation in the left hemisphere; Right, mean cross-correlation in the right hemisphere; Combined, mean cross-correlation across both hemispheres; Combined Max, a reference line drawn at the maximum value of the combined curve. The shadow reflects standard error.

**Supplementary Figure 2**

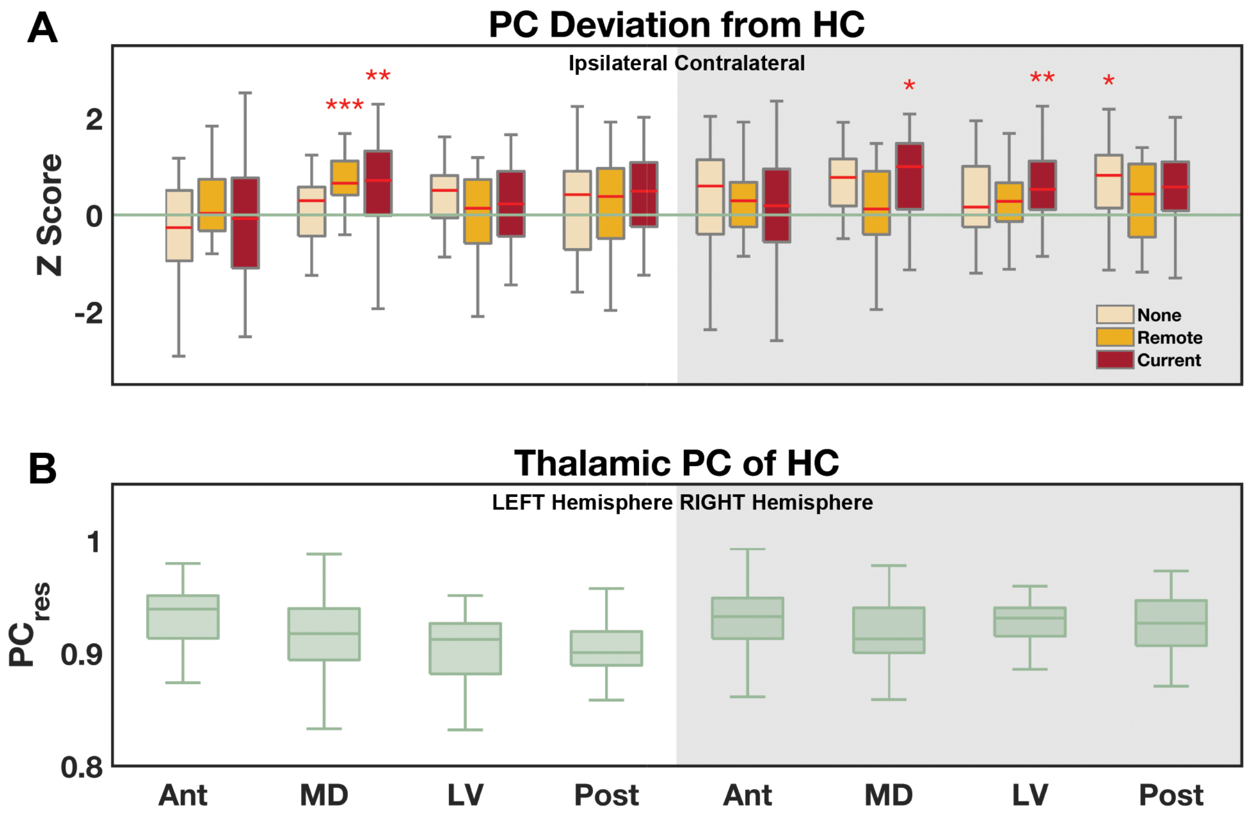

**Supplementary Figure 2. Deviation scores of the thalamic participation coefficient (PC) of the three patient groups.** (A) The deviation score was estimated in reference to data obtained from a matched healthy control group (HC), presented ipsilaterally and contralaterally. The green reference line represents a zero deviation; that is, the mean of the healthy control group. Selective PC increases were observed, and marked by asterisks. (B) Thalamic PC values of the healthy control group are presented in the left and right hemisphere. Res: residual after confound regression; Ant, anterior, MD, medial-dorsal, LV, lateral-ventral, Post, posterior thalamic nuclear groups. *, $P < 0.05$; **, $P < 0.01$; ***, $P < 0.001$. Statistics were obtained via a non-parametric permutation test controlling for multiple comparisons. The central mark indicates the median, and the bottom and top edges of the box indicate the 25$^{th}$ and 75$^{th}$ percentiles, respectively.

**Supplementary Figure 3**

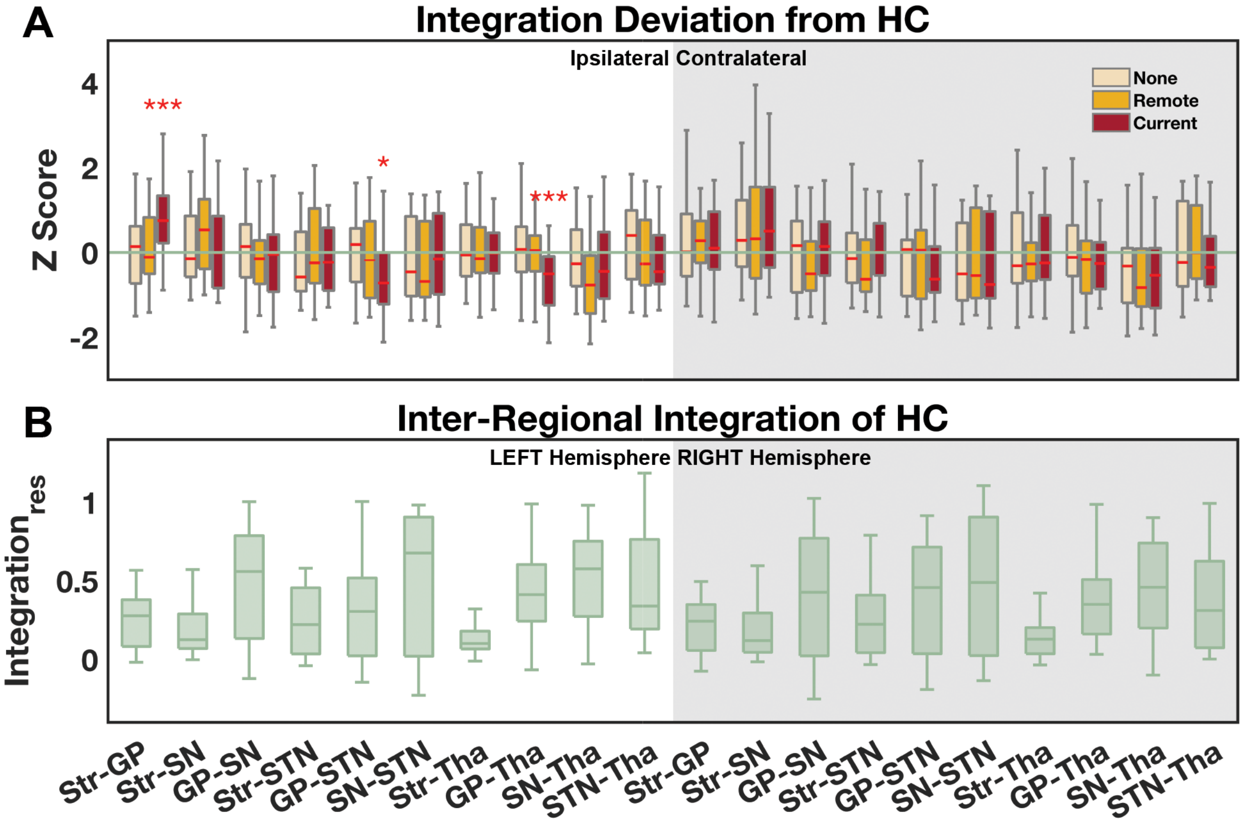

**Supplementary Figure 3. Deviation scores of the pairwise interregional integration of the three patient groups.** (A) The deviation score was estimated in reference to data obtained from a matched healthy control group (HC), presented ipsilaterally and contralaterally. The green reference line represents a zero deviation; that is, the mean of the healthy control group. Selective integration atypicalities (both increases and decreases) were observed, and marked by asterisks. (B) Pairwise interregional integration values of the healthy control group are presented in the left and right hemisphere. Res: residual after confound regression; Str, striatum; GP, globus pallidus; SN, substantia nigra; STN, subthalamic nucleus; Tha, thalamus. *, $P < 0.05$; **, $P < 0.01$; ***, $P < 0.001$. Statistics were obtained via a non-parametric permutation test controlling for multiple comparisons. The central mark indicates the median, and the bottom and top edges of the box indicate the 25$^{th}$ and 75$^{th}$ percentiles, respectively.

**Supplementary Figure 4**

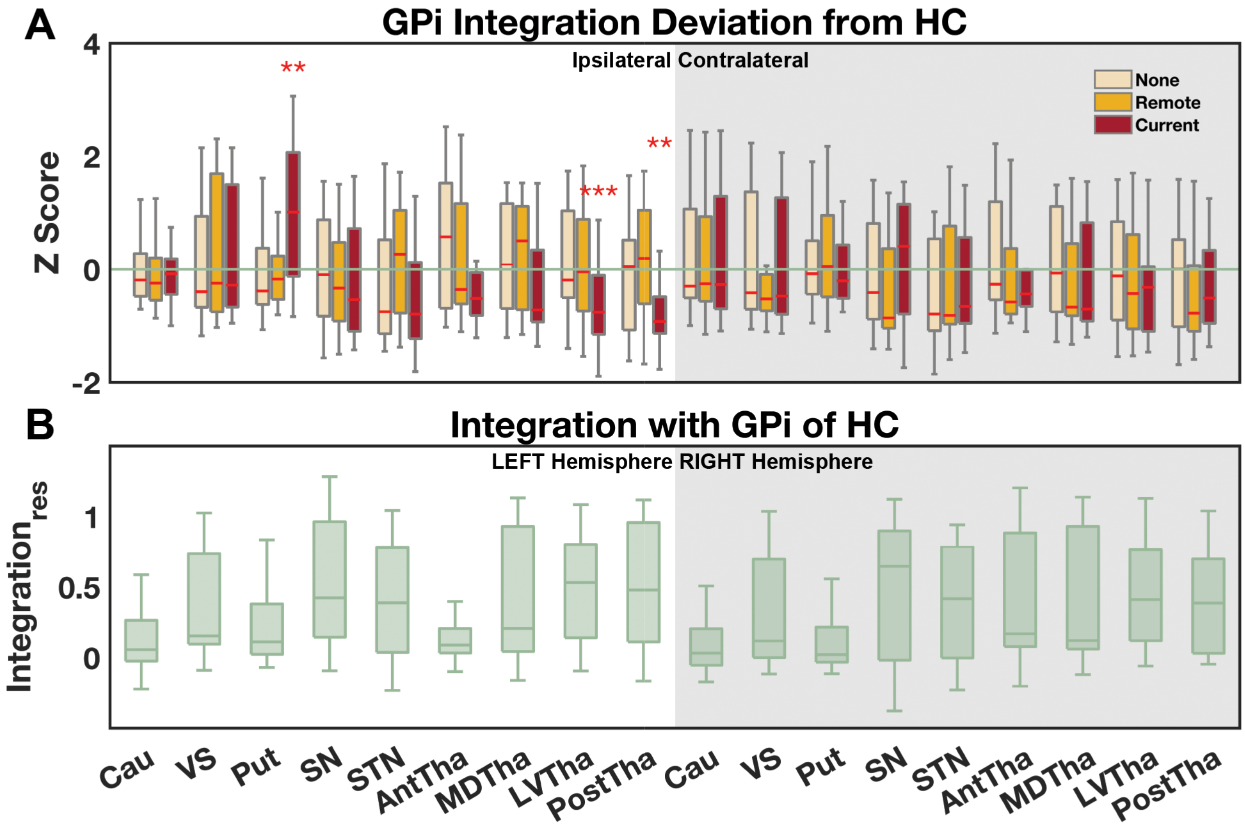

**Supplementary Figure 4. Deviation scores of the interregional integration with GPi of the three patient groups.** (A) The deviation score was estimated in reference to data obtained from a matched healthy control group (HC), presented ipsilaterally and contralaterally. The green reference line represents a zero deviation; that is, the mean of the healthy control group. Selective integration atypicalities (both increases and decreases) were observed, and marked by asterisks. (B) Interregional integration values of the healthy control group are presented in the left and right hemisphere. Res: residual after confound variable regression; GPe, globus pallidus externus; GPi, globus pallidus internus; Cau, caudate; VS, ventral striatum; Put, putamen; SN, substantia nigra; STN, subthalamic nucleus; Tha, thalamus; Ant, anterior, MD, medial-dorsal, LV, lateral-ventral, Post, posterior nuclear groups. *, $P < 0.05$; **, $P < 0.01$; ***, $P < 0.001$. Statistics were obtained via a non-parametric permutation test controlling for multiple comparisons. The central mark indicates the median, and the bottom and top edges of the box indicate the 25$^{th}$ and 75$^{th}$ percentiles, respectively.
